# Genome-wide CRISPR knockout screening with viral replicons for identification of host factors involved in viral replication

**DOI:** 10.1101/2025.01.09.632058

**Authors:** Karen W. Cheng, Madhura Bhave, Andrew Markhard, Duo Peng, Karan D. Bhatt, Katherine Travisano, Josette V. Medicielo, Sanae Lembirik, Leila Njoya, Manu Anantpadma, Jens H. Kuhn, Andreas S. Puschnik, Amy Kistler

## Abstract

Pooled CRISPR knockout (KO) screens using live viruses are a proven and valuable approach for identifying essential host factors acting across viral life cycles. Here we describe the development of a pooled genome-wide CRISPR KO screening approach using stable viral replicon cell lines to specifically identify host factors essential for viral replication. Virus replicons are non-infectious, therefore enabling the study of highly virulent viruses under standard biosafety level 2 containment. We developed a stable fluorescent dengue virus type 2 (DENV-2) replicon cell line to perform a pooled genome-wide FACS-based CRISPR KO screen. This benchmark DENV-2 replicon screen successfully identified host genes previously known to be required for viral DENV-2 replication (e.g., endoplasmic reticulum membrane complex and oligosaccharyltransferase complex components) and confirmed two additional genes (DOHH and ZFP36L2) involved in replication that have not been recovered in prior live virus screens. We applied this replicon screening approach to two highly divergent viruses: chikungunya virus (CHIKV), a positive-sense RNA virus that replicates at the plasma membrane, and Ebola virus (EBOV), a negative-sense RNA virus that replicates in cytoplasmic inclusion bodies. The CHIKV replicon screen identified two genes known to be required for replication (G3BP1 and G3BP2) and several additional, novel genes (CLEC4G, CSDE1, GOLGA7, HNF1A, and PCBD1). We verified two of them (CSDE1 and GOLGA7) in live CHIKV replication assays. A distinct set of genes (EHMT1, EHMT2, and USP7) were recovered in the EBOV replicon screen and were further confirmed using independent transient transfection assays. Thus, viral replicon-based screens provide a useful approach that can be extended to viruses of diverse taxa to identify host pathways essential for viral replication and to uncover potential novel targets for host-directed medical countermeasures.

## Introduction

Elucidating the spectrum of host cell genes and pathways that viruses use over the course of infection is critical to advance our basic understanding of virus biology and pathogenicity, the cellular determinants of host range for emerging/re-emerging viral pathogens, and potential targets for novel host-directed antiviral development^1–5^. The application of genome-wide CRISPR knockout (KO) screens to live virus infections has served as a powerful strategy to comprehensively identify host genes and cellular pathways that are essential for infection across a wide range of diverse viruses. For example, within a year of the emergence of severe acute respiratory syndrome coronavirus 2 (SARS-CoV-2), the set of overlapping and non-overlapping host genes and pathways required for SARS-CoV-2 was broadly delineated through a series of independent genome-wide CRISPR ^6^. Likewise, live virus CRISPR KO screens with dengue virus type 2 (DENV-2) have yielded a suite of novel genes and pathways that were independently shown to play a functional role in viral entry as well as the formation and function of the DENV replication complex during infection ^7–10^.

However, pooled CRISPR screens with live viruses are quite variable regarding identification of host factor dependencies spanning entire viral life cycles ^3^. This is, in part, a reflection of the basic format of live virus CRISPR KO screens that typically entail selecting for cells harboring KOs that confer survival during infections. Because viral entry occurs upstream of all subsequent intracellular stages of infection (viral transcription, translation, replication, assembly, and egress), the loss of host genes/pathways required for entry would be expected to dominate the genes and host pathways enriched in these screens. Likewise, the biology of a virus also plays a role (e.g., cellular receptor usage patterns, kinetics of viral translation, transcription, replication, and egress). Nonetheless, there are notable exceptions: genome-wide CRISPR KO screens with live viruses have successfully uncovered cellular genes/pathways for steps downstream of entry (e.g., viruses with multiple cellular receptors, including DENV-2 ^7–10^, West Nile virus ^11,12^, hepatitis A virus ^13^, and several enteroviruses ^14^). Given these limitations, alternative CRISPR screening modalities are needed to improve our ability to identify the host factor dependencies essential for different stages of the viral life cycle.

To date, multiple variations in CRISPR screening approaches have been developed to try to specifically identify host genes/pathways required for stages of the viral life cycle that are downstream of entry. For example, a recent FACS-based CRISPR KO screen with yellow fever virus encoding an ectopically expressed fluorescent protein was developed to specifically identify cellular genes required for efficient translation of viral proteins ^14^. In a completely different approach, libraries of guide RNAs (gRNAs) targeting cellular host genes were encoded within the genomes of human influenza A virus (FLUAV) and human cytomegalovirus (HCMV) and then used to launch a screen to identify host genes that, when activated, enhance or suppress viral production over multiple rounds of infection ^15,16^. Each of these approaches has identified novel host genes/pathways acting downstream of entry. However, we still lack a screening approach that specifically identifies host genes required for viral replication.

Viral replicons are sub-genomic virus surrogate systems consisting of virus genomes in which the genes encoding the structural proteins required for virion formation are replaced with reporter or reporter-selection gene cassettes ^17,18^. Introduction of viral replicon RNA, which encodes all the non-structural genes harboring the enzymatic activities required for viral replication and transcription, into cells can confer autonomous replication of the replicon. Thus, for instance, fluorescence activity/drug resistance may serve as a reporter for viral transcription, translation and replication. Importantly, viral replicons bypass both the upstream entry and downstream egress phases of the viral life cycle. Together, these two features offer the added benefit of providing a non-infectious assay system to investigate replication requirements for any virus of interest in the context of a biosafety level 2 (BSL-2) environment.

We explored the feasibility of genome-wide CRISPR KO screens with stable viral replicon cell lines to specifically recover host factors required for viral replication in three distinct RNA viruses: two positive-sense RNA viruses (DENV-2 and chikungunya virus [CHIKV]) and a negative-sense RNA virus (Ebola virus [EBOV]). We successfully performed genome-wide CRISPR KO screens with replicons for all three of the viruses and identified host factors involved in their replication. We further verified the hits from our CRISPR KO screen in a live virus context for DENV and CHIKV and independent transient replicon assays for EBOV. Together, our data indicate that viral replicons can serve as an important tool to study replication of diverse classes of even highly virulent viruses in BSL-2 facilities.

## Results

Generation of a stable and responsive DENV-2 replicon reporter cell line suitable for genome-wide CRISPR KO screening To benchmark the performance of a genome-wide CRISPR replicon screening approach, we first focused on DENV-2, for which host factors involved in viral replication have been well-described through both targeted experiments and multiple live-virus CRISPR screens ^3^.

Generation of a viral replicon cell line that exhibits stable and homogenous reporter activity and is responsive to inhibition is a prerequisite for pooled replicon-based CRISPR screening. We created a DENV-2 replicon cell line that could confer stable reporter activity by first replacing the viral genomic region encoding the structural proteins in the DENV-2 16681 infectious clone with a reporter-selection cassette encoding enhanced green fluorescent protein (eGFP) fused to a selectable marker (Supplemental Figure 1A and 1B). We generated two distinct reporter-selection cassettes (“eGFP-blasticidin” and “eGFP-Zeocin”; Supplemental Figure 1B) to test in the replicon, because the choice of selectable marker can affect the levels and heterogeneity of reporter protein expression ^19^. Each reporter-selection cassette was assayed for its ability to maintain viral replication over several weeks—after RNA electroporation and initial antibiotic selection (Figure 1A, 1B). Human embryonic kidney (HEK) 293T cells transfected with the eGFP-blasticidin DENV-2 replicon RNA were enriched as measured by eGFP expression; however, the percentage of eGFP^+^ cells plateaued and decreased over time despite increasing blasticidin concentration (Figure 1B). In contrast, HEK 293T cells transfected with the eGFP-Zeocin replicon cell line were enriched to a homogenous and stable eGFP^+^ cell population (Figure 1B). The fluorescence signal of the eGFP-Zeocin replicon cell line correlated with expression of DENV-2 nonstructural protein NS3, a key component of the DENV-2 replication complex, and with double-stranded RNA (dsRNA), indicating the eGFP reporter activity in the eGFP-Zeocin replicon cell line reflected the presence and replication of the DENV-2 replicon RNA (Pearson’s correlation coefficient = 0.78 and 0.45, for eGFP/NS3 and eGFP/dsRNA correlations, respectively; Figure 1C).

**Figure 1.**
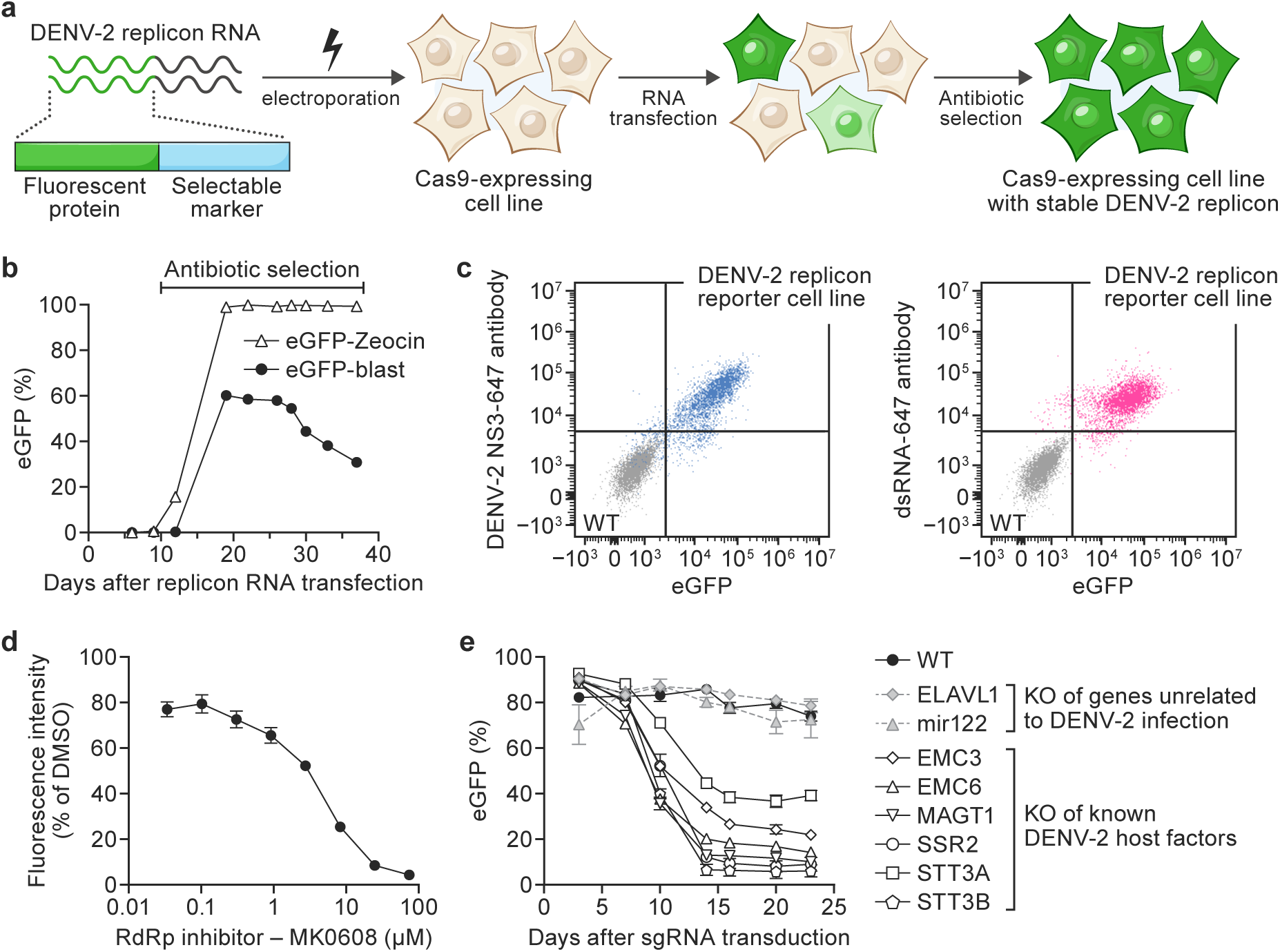
Generation of a stable and responsive dengue virus type 2 replicon reporter cell line suitable for genome-wide CRISPR knockout screening. A, Overview of DENV-2 stable replicon reporter cell line generation: Cas9-expressing cells were transfected with a DENV-2 replicon RNA transcribed *in vitro*. Cells that receive the replicon RNA and successfully replicate the encoded viral non-structural genes and enhanced green fluorescent protein (eGFP) reporter-selection cassette begin to fluoresce. Growth in the presence of antibiotics selects a stable DENV-2 replicon cell line with a homogenous fluorescent signal. B, Time-course of eGFP expression measured by flow cytometry in human embryonic kidney (HEK) 293T-Cas9 cells transfected with DENV-2 replicon RNA with different reporter cassette variants (eGFP-blasticidin vs eGFP-Zeocin) grown in the presence of blasticidin or Zeocin. C, Flow cytometry analysis of the correlation of fluorescent signal from the DENV-2 replicon eGFP reporter and from DENV-2 NS3 antibody staining (left panel) or dsRNA antibody staining (right panel) in HEK293T-Cas9 (gray) and the stable HEK293T-Cas9 DENV-2 replicon cell line (blue or magenta). D, Pharmacological inhibition of the stable Huh7.5.1-Cas9 DENV-2 replicon cell line with an RNA-dependent RNA polymerase (RdRp) inhibitor (MK608) or control (dimethylsulfoxide [DMSO]). Experiment was performed in technical triplicate and values represent mean fluorescence intensity ± SD relative to DMSO control. E, Inhibition of known DENV-2 host factors by CRISPR KO in the stable Huh7.5.1-Cas9 DENV-2 replicon cell line transduced with lentiviruses harboring the guide RNAs (gRNAs) indicated in the plot legend. DENV-2 replicon activity was quantified as changes in % eGFP reporter expression upon CRISPR knockout (KO) of host genes established to be involved in DENV-2 replication (open symbols) or genes unrelated to DENV-2 replication (gray symbols). Wild-type (WT) DENV-2 replicon cells were assayed in parallel (black symbols). Experiments were performed in technical triplicates and values represent mean % eGFP ± SD expression level normalized to % eGFP at the start of the experiment. Gene names are abbreviated according to the human standard (https://www.genenames.org/).

Next, we rationalized that responsiveness of the eGFP signal to perturbations of viral replication is critical. To improve the system, we appended a PEST sequence, known to reduce the intracellular half-life of proteins ^20^, to the marker cassette (“Zeocin-eGFP-PEST”; Supplemental Figure 1B). The optimized DENV-2 replicon was stably transfected into Huh7.5.1–Cas9 cells, an established cell line model for studying DENV-2 infection and previously used for DENV-2 host factor screens ^8^ (Supplemental Figure 1B). Western blot analysis confirmed expression of the DENV-2 nonstructural proteins NS2B, NS3, and NS4B and the eGFP reporter (Supplemental Figure 1C). To test whether the Huh7.5.1–Cas9 DENV-2 replicon cell line (hereafter referred to as DENV-2 replicon cell line) was suitable for screening assays, we assessed its responsiveness to 7-deaza-2’-C-methyladenosine (MK-0608), a known RNA-directed RNA polymerase inhibitor of DENV-2 replication ^21^. As expected, MK-0608 decreased the eGFP reporter signal in a dose-dependent manner (Figure 1D). We next evaluated the impact of CRISPR KO of genes encoding components of the endoplasmic reticulum (ER) membrane protein complex (EMC) and oligosaccharyltransferase (OST) complexes that have been established host factor complexes required for DENV-2 replication and protein expression ^8,22^. These KOs resulted in a decreased eGFP reporter signal in the DENV-2 replicon cell line, whereas CRISPR KO of ELAVL1, a gene known not to affect DENV-2 replication, had no effect (Figure 1E). Together, these experiments demonstrated that the eGFP signal in the DENV-2 replicon cell line was both stable and responsive to replication inhibition and thus suitable for pooled genome-wide CRISPR screening.

### CRISPR replicon screen identifies known and novel host factors involved in DENV-2 translation and replication

We next performed a pooled, genome-wide CRISPR KO screen with the Human Brunello pooled sgRNA library ^23^ in the DENV-2 replicon cell line to benchmark the viral replicon screening approach. Using a FACS-based readout, we isolated a cell population in which the eGFP signal had decreased over the course of the screen (“eGFP-low”), likely corresponding to CRISPR KO of host genes important for the DENV-2 replicon activity, as well as a control “eGFP-high” sorted cell population (Figure 2A). The cells were then analyzed by high-throughput sequencing of gRNA abundances and bioinformatic analysis using MAGeCK software ^24^.

**Figure 2.**
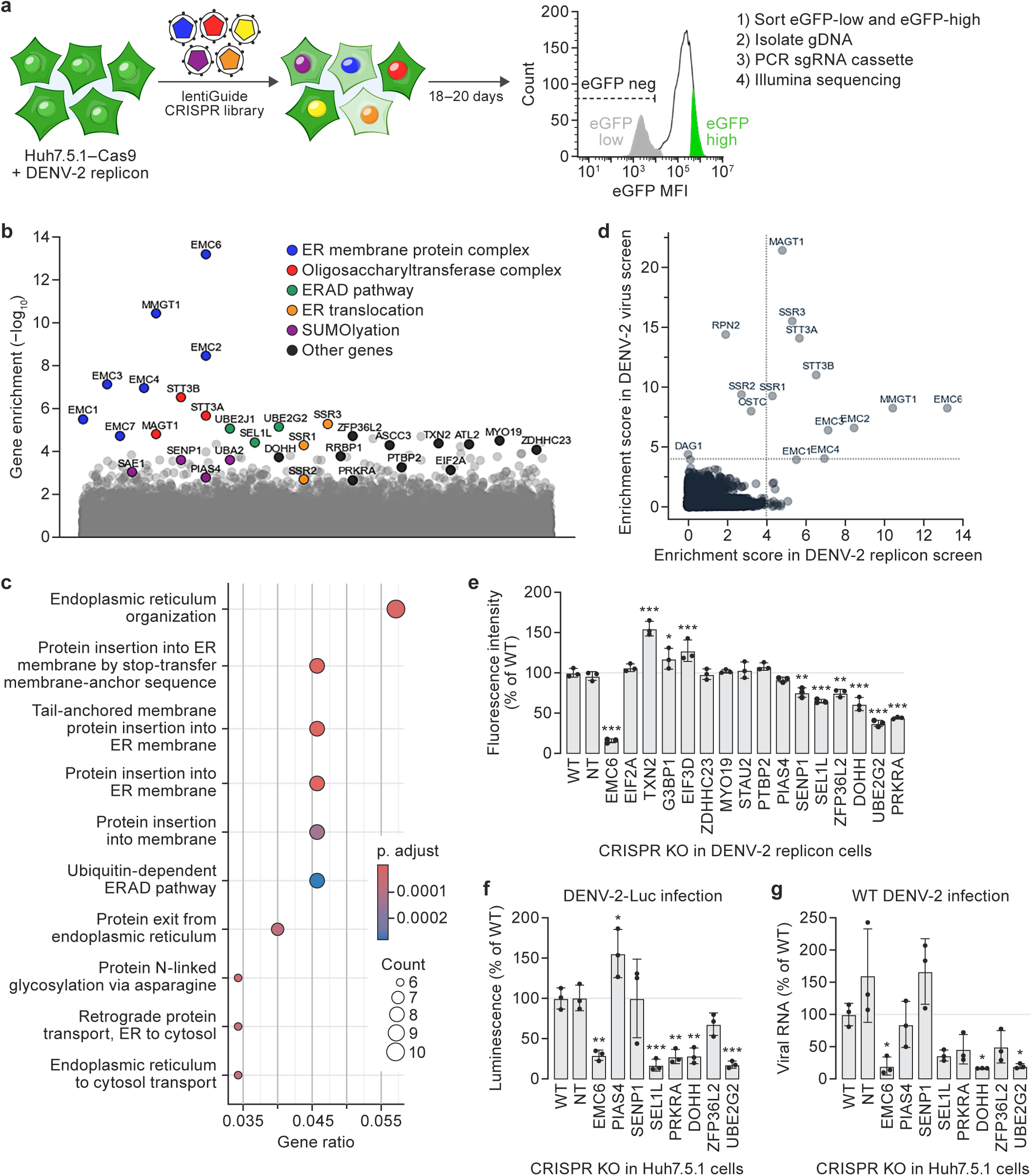
CRISPR replicon screen identifies known and novel host factors involved in dengue virus type 2 translation and replication. A, Schematic of the genome-wide CRISPR knockout (KO) screening approach. B, Enrichment plot from CRISPR KO screen of dengue virus type 2 (DENV-2) replicon cell line for host factors involved in DENV-2 replication. The y-axis represents the significance of enrichment calculated from MAGeCK RRA statistical analysis of genes that were enriched in the selected (eGFP-low) population vs control (eGFP-high) cell population; the x-axis corresponds to identified genes. C, Pathway enrichment summary plot. D, Comparison of hits from genome-wide CRISPR KO screens with replicon (x-axis, data from this study) versus live virus CRISPR KO screen hits (y-axis, data from (Marceau et al., 2016)). Dotted lines are the cutoffs for false discovery rate (FDR) <10%. The upper right quadrant represents overlapping hits, the upper left quadrant represents genes enriched only in the live virus screen, and the lower right quadrant represents genes enriched only in the replicon screen. E, Independent replicon screen phenotype validation. Arrayed CRISPR KO in the DENV-2 replicon cell line targeting a panel of candidate genes identified in the screen. eGFP fluorescence intensity 18 d after cells were treated with individual targeting gRNAs (or non-targeting control gRNA, NT) was measured to quantify DENV-2 replicon activity. Values represent mean fluorescence intensity ± SD normalized to wild-type (WT) control from two biological replicates. F, Quantification of KO impact on live DENV-2 infections. Parental (WT) Huh7.5.1 and independently generated KO cell populations were exposed to DENV-2-Luc (multiplicity of infection [MOI] = 0.1) and harvested 48 h after exposure. DENV-2 infection efficiency in each condition was calculated as a ratio of luminescence (DENV-2-Luc reporter expression) to Hoechst stain (proxy for cell count) and normalized to WT. Experiments were performed in technical triplicates and values represent mean % eGFP ± SD. G, RT-qPCR of DENV RNA in Huh7.5.1. KO cell lines infected with DENV-2 16681 (MOI = 0.1) and harvested 48 h post-infection. Values represent mean viral RNA expression ± SD normalized to WT control from three biological replicates. Gene names are abbreviated according to the human standard (https://www.genenames.org/).

Among the top enriched genes recovered in the DENV-2 replicon screen were previously identified host genes that are important for viral genome replication or biogenesis and stability of viral proteins. These genes included components of the EMC and OST complexes, the ER-associated degradation (ERAD) pathway, and the ER translocon (Figure 2B and 2C). Previously, additionally identified RNA-binding proteins involved in the replication of the viral RNA (vRNA) genome (such as, ASCC3 and RRBP1) were also enriched ^10^. To comprehensively assess the extent to which the replicon screen recovered host genes known to be involved in DENV-2 replication, we compared the full gene enrichment datasets of the replicon screen and a previously published genome-wide CRISPR KO survival screen performed in Huh7.5.1 cells infected with live DENV-2 ^8^. Components corresponding to key host complexes required for DENV-2 replication complex and formation were among the most significantly enriched genes in both the replicon and live virus screens (Figure 2D).

We next investigated whether the replicon-based CRISPR screening approach could identify host factors required for DENV-2 replicon activity that had not been identified in previous survival screens with DENV-2 infection. We compared the top 200 hits (top 1%) from our replicon screen to other live DENV-2 survival CRISPR screens ^7,8,10^ and selected a subset of genes for further validation that were unique to this replicon screen: DOHH, EIF2A, EIF3D, G3BP1, MYO19, PIAS4, PRKRA, PTBP2, SEL1L, SENP1, STAU2, TXN2, UBE2G2, ZDHHC23, and ZFP36L2 (Figure 2E). The recovery of SEL1L and UBE2G2 among this set are notable. Although not recovered in prior genome-wide DENV-2 CRISPR KO screens, SEL1L was identified to be involved in DENV-2 in a separate insertional mutagenesis screen in human haploid cells ^8^. Likewise, although UBE2G2 had not been specifically identified in previous survival screens for DENV-2, it is involved in the ERAD pathway and has been previously implicated in genome-wide CRISPR KO screens with West Nile virus, a virus closely related to DENV-2, to modulate orthoflavivirus protein homeostasis during infections ^25,26^. The specific enrichment of these and the other additional factors in the replicon screen highlights the potential of the replicon screen to identify distinct host factors involved in the viral life cycle.

To independently confirm whether these genes were important for DENV-2 replication, we knocked out each gene individually in the parental DENV-2 replicon cell line using the top-performing gRNAs from the Brunello library for each hit from the replicon screen. Genotyping analysis confirmed each KO cell pool (Supplemental Figure 2A), and flow cytometry measured the changes in replicon reporter fluorescence signal. Of the 15 candidate genes selected for verification, DOHH, SEL1L, SENP1, PRKRA, UBE2G2, and ZFP36L2 KO significantly decreased in fluorescence intensity relative to that in a control cell line transduced with a non-targeting gRNA (Figure 2E). To determine whether these candidate hits were also relevant in a live DENV-2 infection context, we generated parallel KOs in the parental Huh7.5.1 cell lines and then exposed the resulting cell lines to either a DENV-2-luciferase reporter virus (DENV-2-Luc) or wild-type DENV-2 at a low (0.1) multiplicity of infection for 48 h. These analyses showed that depletion of DOHH, PRKRA, SEL1L, UBE2G2, and ZFP36L2 consistently decreased DENV-2 infection of Huh7.5.1 cells both at the level of viral protein translation (Figure 2E) and RNA replication (Figure 2F). Taken together, these data demonstrate that genome-wide CRISPR screens with stable viral replicon cell lines can be used to discover both known and novel host factors involved in viral replication and translation processes.

### Genome-wide CRISPR KO screen with a stable CHIKV replicon cell line

We next tested whether the CRISPR replicon screening approach could identify host factors involved in the replication of another positive-sense RNA virus, CHIKV. CHIKV shares similarities with the DENV-2 life cycle—such as, initial RNA release in the cytoplasm, translation, and polyprotein processing—but CHIKV belongs to a different viral family than DENV-2 (*Togaviridae* vs. *Flaviviridae*) and has distinct genome and replication features ^25^. Notably, CHIKV has not one but two open reading frames (ORFs): one for nonstructural proteins and another for structural proteins, transcribed from a sub-genomic promoter. Additionally, its replication complex functions at the plasma membrane in invaginated spherules^27^. Despite recent advances in understanding CHIKV’s replicase complex architecture at high resolution ^28^, to date, live CHIKV CRISPR screens have only provided detailed understanding of the cellular receptor, MXRA8 ^29^, and the role of a single gene involved in CHIKV replication, FLH1 ^30^.

We adapted the DENV-2 replicon screening method for CHIKV by modifying an established CHIKV replicon ^31^. We tested several Zeocin-eGFP reporter-selection cassette variants in this system and found that the eGFP-Zeocin variant exhibited stable fluorescence and detectable expression of CHIKV non-structural proteins and eGFP (Supplemental Figures 3A and 3B). In this CHIKV replicon cell line, depletion of G3BP1, a known CHIKV replication factor identified through prior targeted studies ^32^, decreased the fraction of fluorescent cells, whereas knockout of the CHIKV cellular receptor MXRA8 ^29^ had no effect on replicon reporter activity relative to that observed in wild-type cells (Figure 3B). Thus, the CHIKV replicon cell line was sufficiently stable and responsive to perturbation for pooled CRISPR screening.

**Figure 3.**
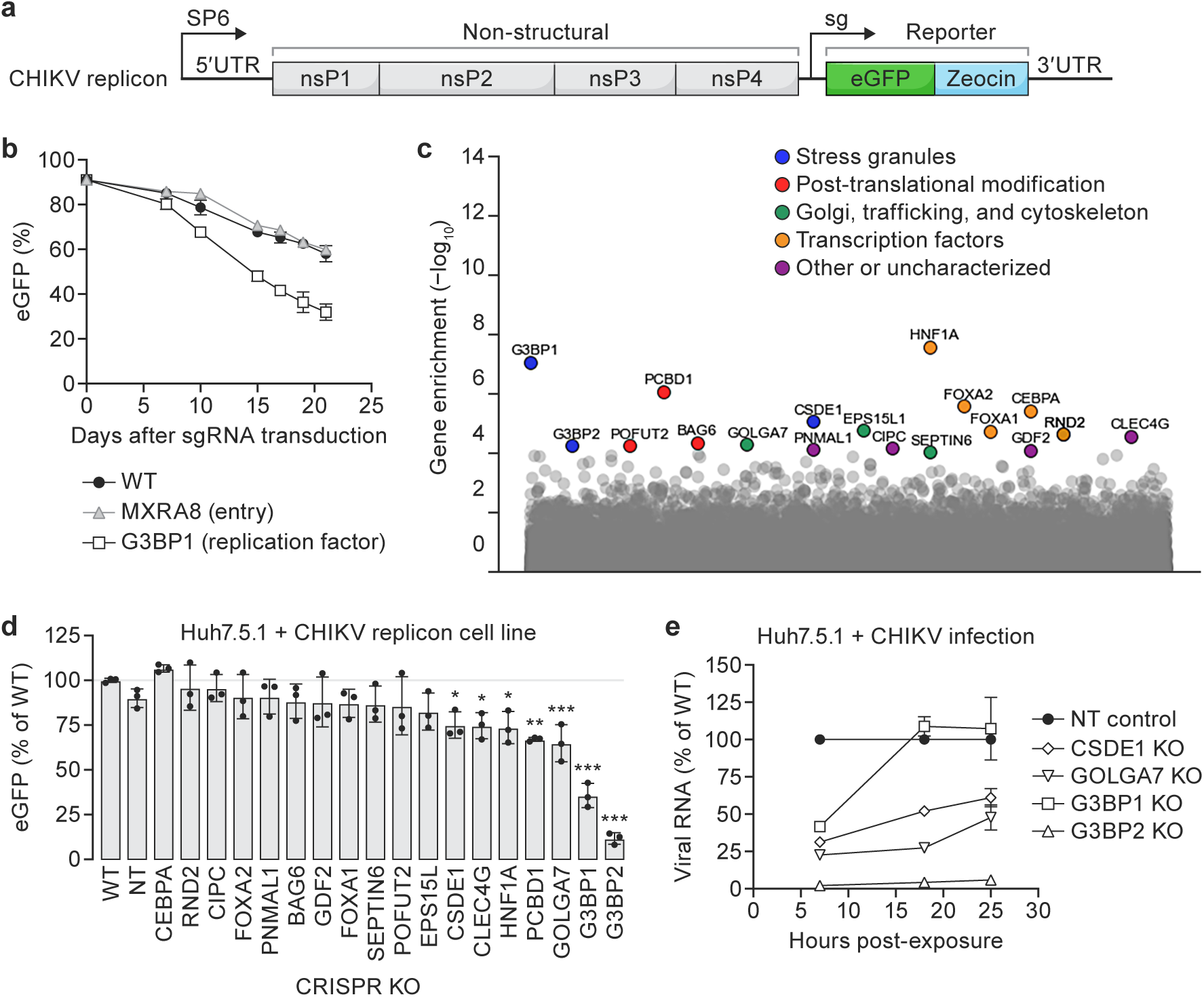
Genome-wide CRISPR knockout screen with a stable chikungunya virus replicon cell line. A, Schematic of chikungunya virus (CHIKV) replicon used in this study. B, Inhibition of host factors known to be required for CHIKV replication by CRISPR knockout (KO) in the stable Huh7.5.1-Cas9 CHIKV replicon cell line. CHIKV replication levels were quantified by changes in % eGFP expression upon CRISPR KO of host genes established to be involved in CHIKV replication (open symbols) or genes unrelated to CHIKV replication (gray), with wild-type (WT) CHIKV replicon cells assayed in parallel (black). Experiments were performed in technical triplicates and values represent mean % eGFP ± SD normalized to % eGFP at the start of the experiment. C, Enrichment plot from a CRISPR KO screen with the CHIKV replicon cell line for host factors involved in CHIKV replication. The y-axis represents the significance of enrichment calculated by MAGeCK RRA statistical analysis of genes that were enriched in the selected (eGFP-low) population vs control (unselected) cell population; the x-axis corresponds to genes. D, Independent validation of replicon screen phenotypes: arrayed CRISPR KO in the CHIKV replicon cell line targeting a panel of candidate genes identified in the screen. The fraction of eGFP^+^ cells 18 d after cells were treated with individual targeting guide RNA gRNAs or (non-targeting control gRNA, NT) was measured to quantify CHIKV replicon activity. Values represent mean % eGFP ± SD normalized to WT control from 3 biological replicates. E, Quantification of KO impact on live CHIKV infections. Parental (WT) Huh7.5.1 and independently generated KO cell populations were exposed with CHIKV LR-2006 OPY1 (MOI 0.1) and harvested at 7, 18, and 25 h after exposure. Values for each timepoint represent the mean ± SD of normalized CHIKV expression levels relative to the normalized levels detected in the control cell lines transduced with non-targeting gRNA. Gene names are abbreviated according to the human standard (https://www.genenames.org/).

To identify host factors required for CHIKV replication, we applied the pooled genome-wide CRISPR KO replicon screening approach established for DENV-2. We performed two replicate screens using the Brunello library and the same FACS-based readout for CHIKV replicon activity. An eGFP-low population of cells were sorted and collected at the end of the screen, and high-throughput sequencing and bioinformatic analysis were performed to determine the enrichment of depleted genes in the screen based on the abundance of gRNAs in the sorted eGFP-low population compared to an unselected control population of CHIKV replicon cells transduced with the gRNA library and grown in parallel during the screen ^24^.

A set of 18 significantly enriched genes were identified in the CHIKV replicon screen, with several types of functional classes represented among them (Figure 3C). Genes involved in stress granule biogenesis (CSDE1, G3BP1, and G3BP2) ranked among the top hits recovered in the screen. Additionally, genes implicated in Golgi trafficking (EPS15L1, GOLGA7, and SEPTIN6), post-translational modifications (BAG6, PCBD1, and POFUT2), and transcription regulation (CEBPA, FOXA1, FOXA2, HNF1A, and RND2) were also identified. A fifth group of enriched genes mapped across a diverse set of potential functional classes (CIPC, CLEC4G, GDF2, and PNMAL). Individual genes were knocked out in the CHIKV replicon cell line for independent validation of their impact on CHIKV replicon activity. As expected, depletion of G3BP1 and G3BP2 genes were significantly reduced in eGFP^+^ cells relative to untreated CHIKV replicon cells or CHIKV replicon cells transduced with a non-targeting gRNA. The phenotype of five additional genes enriched in the screen were similarly verified: CLEC4G, CSDE1, GOLGA7, HNF1A, and PCBD1 (Figure 3D).

Of these verified hits, CSDE1 and GOLGA7 were of particular interest for immediate follow-up studies based on their potential links to the CHIKV replication cycle and subcellular localizations. Cold shock domain containing E1 (CSDE1) is an RNA-binding protein implicated in diverse aspects of RNA biology (translation and catabolism) and regulation of stress granule formation ^33,34^; it is known to be used by CHIKV non-structural protein 3 (nsP3) during infection and required for viral replication ^31,32,35,36^. CSDE1 localization within the cell also overlaps the compartments that CHIKV transits during its life cycle (Golgi apparatus, cytoplasm, and plasma membrane). Golgin A7 (GOLGA7) is a component of the Golgi membrane involved in trafficking between the Golgi apparatus and plasma membrane ^37^, a potentially important process relevant for the formation of CHIKV replication compartment.

We investigated the impact of depleting CSDE1, G3BP1, G3BP2, and GOLGA7 in Huh7.5.1 cells during CHIKV infections. We generated polyclonal populations of a control cell line harboring a non-targeting gRNA (NT) and KO cells for each of the gene targets of interest (Supplemental Figure 3C). We infected these cells with a CHIKV isolate that is isogenic to the replicon (LR-2006 OPY1) and measured relative levels of vRNA present in the KO pools compared to the non-targeting control cell line (Figure 3E).

At the earliest timepoint (7 h), we observed a decrease in the levels of CHIKV RNA in all KO cell lines relative to the NT cell line (Figure 3E). Subsequent timepoints revealed distinct viral replication phenotypes depending on the depleted gene. The CHIKV replication phenotypes in the G3BP1 and G3BP2 KO cell lines were consistent with previously published observations that depletion of G3BP1 modestly decreases CHIKV replication whereas depletion of G3BP2 has a much stronger effect ^32^. For the CSDE1 KO and GOLGA7 KO cells, an intermediate CHIKV replication phenotype was observed: CHIKV RNA abundance increased over the course of the infection time course, but it remained well below the levels observed in the NT control cells. Full analysis of all verified hits recovered from the screen is still in progress; however, these data confirm that the genome-wide CRISPR screen with the CHIKV replicon targeted host factors involved in replication.

### Genome-wide CRISPR KO screen with an EBOV replicon cell line

To assess the extensibility of the CRISPR replicon screening approach, we next applied it to EBOV, a highly virulent negative-sense RNA virus (Supplemental Figure 4A) that has a distinct biology and type of replicon system compared to DENV-2 and CHIKV ^38^. To develop a stable EBOV replicon cell line suitable for FACS-based pooled CRISPR screening under BSL-2 containment, we adapted previously described methods to generate stable EBOV replicon cell lines ^39,40^ (Figure 4A). We first integrated a construct encoding a multicistronic transcript ^39^ encoding an uninterrupted ORF of four EBOV genes that are essential for viral replication and transcription (nucleoprotein [NP], which binds and protects the EBOV RNA; two co-factors of the EBOV RNA-dependent RNA polymerase, the 35kDa viral protein [VP35] and the 30kDa viral protein [VP30]; and the large protein [L protein], which encodes the EBOV RNA-dependent RNA polymerase all the other relevant enzymatic activities) into the genome of Huh7.5.1–Cas9 cell line (“4cis”, top panel, Figure 4A; Methods) to create the 4cis cell line. We next transfected the 4cis cell line with an *in vitro* transcribed “minigenome” EBOV RNA template consisting of the EBOV genomic 5ʹ untranslated region (UTR) and 3ʹ UTR flanking a Zeocin-eGFP reporter-selection cassette in the negative-sense orientation (Supplemental Figure 4B). In this system, constitutively expressed EBOV non-structural proteins recognize, replicate, and transcribe the minigenome RNA, conferring eGFP fluorescence and resistance to Zeocin. Supplementation of media with increasing concentrations of Zeocin enabled selection and expansion of a stable population of EBOV replicon cells with >90% eGFP^+^ signal (bottom panel, Figure 4A).

**Figure 4.**
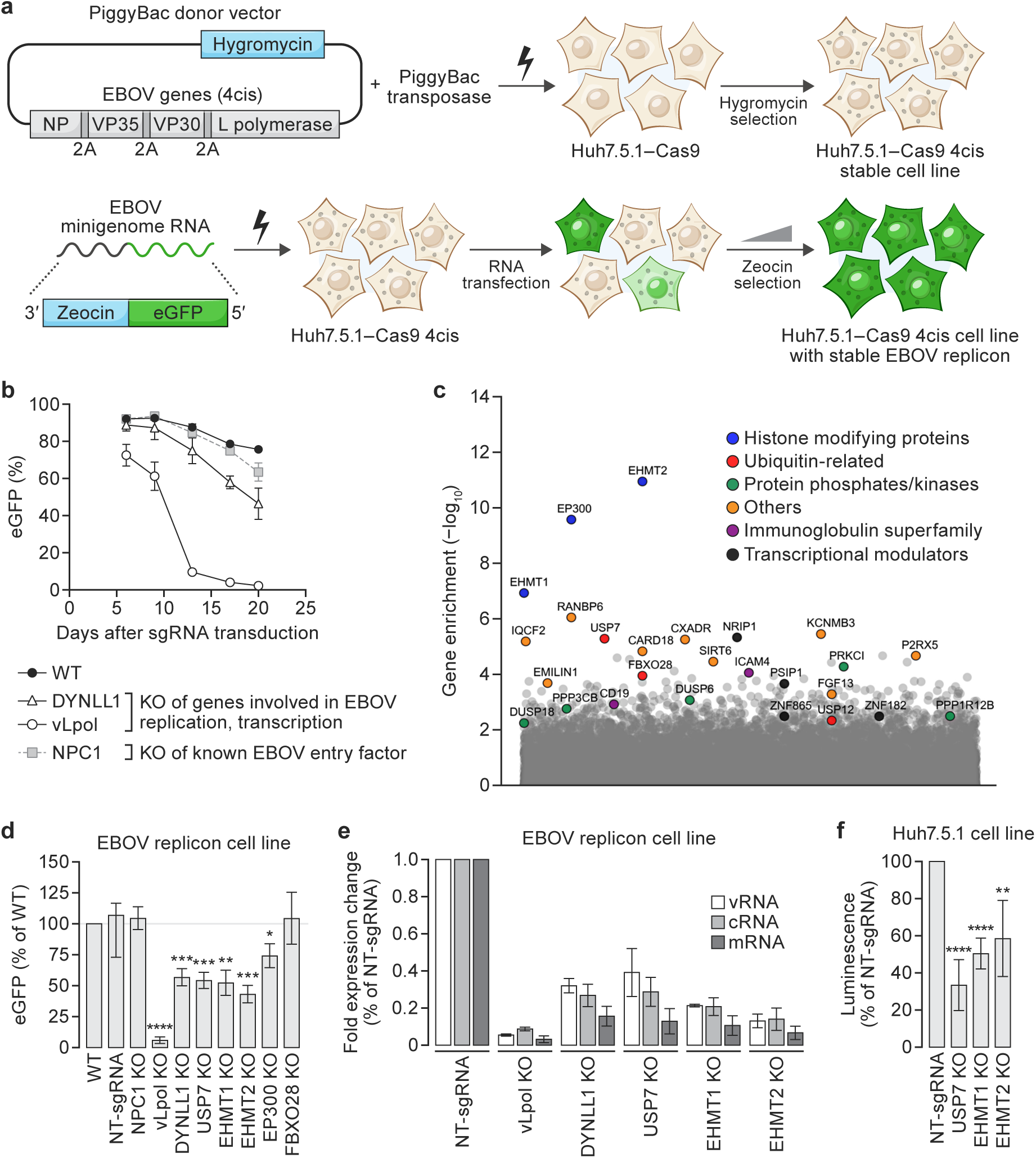
Genome-wide CRISPR knockout screen with an Ebola virus replicon cell line. A, Graphical overview of the steps to generate a stable Ebola virus (EBOV) replicon cell line. PiggyBac transposition was applied to generate a “4cis” cell line, in which four EBOV proteins essential for viral replication and transcription (NP, VP35, VP30 and L protein) are stably expressed after integration and subsequent selection in 100 ug/mL hygromycin media. Next, an *in vitro* transcribed EBOV minigenome RNA (in negative-sense orientation), which encodes the 5ʹ and 3ʹ viral UTRs flanking a reporter-selection (Zeocin-eGFP fusion) cassette, is transfected into the 4cis cell line. Stable population of >90% eGFP^+^ Huh7.5.1-Cas9 EBOV replicon cells were selected and expanded in Zeocin media. B, Validation of the responsiveness of the replicon cell line by CRISPR knockout (KO) of viral L protein and known host factors involved in EBOV entry (Niemann-Pick disease, type C1 [NPC1]) or replication and transcription (dynein light chain LC8-type 1 [DYNLL1]). C, Enrichment plot showing host factor genes recovered from the screen. Enrichment scores are represented as -log10 of the RRA scores obtained from MAGeCK analysis. D, Validation of host factors identified from the screen. Independent KOs of the hits identified in the screen were generated in the EBOV replicon cell line in an arrayed format and flow cytometry was used to measure the eGFP signal over the course of 20 d. The fraction of eGFP^+^ cells on Day 20 of the time course is plotted as mean ± SD (three independent technical replicates). E, Quantification of the levels of different types of EBOV minigenome RNA in the replicon cells harboring different host factor knockouts. Total RNA was isolated from replicon cells with different host factor knockouts and used as input for strand-specific RT-qPCR assay. Plotted values are fold expression changes when normalized to the cells expressing non-targeting gRNA (three independent technical replicates). F, Validation of phenotypes in transient EBOV replicon assays in independently generated KOs in parental cell lines. Effect of host factor knock outs on EBOV replication and transcription in a transient replicon assay with a minigenome encoding a secreted nano-luciferase (SecNLuc). Host factor knockout cell lines and controls were co-transfected with an EBOV 4cis plasmid encoding EBOV NP, VP35, VP30, and L protein, and an EBOV minigenome luciferase reporter plasmid. The SecNLuc signal was measured 48 h post-transfection. Data plotted here represent the mean ± SD (three independent biological replicates). Gene names are abbreviated according to the human standard (https://www.genenames.org/).

Expression and correct processing of EBOV NP, VP35, and VP30 in the EBOV replicon cells was confirmed via western blot analysis (Supplemental Figure 4C and 4D). The eGFP fluorescence detected in EBOV replicon cells served as a reporter for expression of intact functional EBOV L protein. Strand-specific real-time reverse transcription quantitative polymerase chain reaction (RT-qPCR) assay of the EBOV minigenome RNA ^40^ confirmed that all three expected products of L protein transcription and replication activity (vRNA, cRNA, and mRNA) were expressed in the EBOV replicon cells. This analysis showed higher levels of normalized vRNA expression in the EBOV replicon cell line compared to quantitation in the previously described stable EBOV replicon cell line ^40^, but the overall pattern of relative expression levels of each RNA species was the same (vRNA>mRNA>cRNA; Supplemental Figure 4E).

To assess the suitability of EBOV replicon cells for pooled CRISPR screening, we tested whether genetic depletion of EBOV L protein or the dynein light chain LC8-type 1 (DYNLL1) gene, a host factor implicated in EBOV replication ^41^, reduced eGFP reporter activity (Figure 4B). As expected, knockout of EBOV L protein significantly decreased eGFP^+^ cells by Day 5 post-transduction. Similarly, knockout of DYNLL1 reduced eGFP^+^ cells by Day 13 post-transduction (Figure 4B). In contrast, depletion of the Niemann-Pick disease, type C1 gene (NPC1), encoding the EBOV intracellular receptor ^42^, had no significant effect on the fraction of eGFP^+^ cells. These results confirm that the EBOV replicon cell line meets key criteria for pooled CRISPR screening: 1) stable expression of EBOV proteins that drive stable eGFP^+^ signal via replication of the Zeocin-eGFP minigenome and 2) an eGFP signal that responds to depletion of essential viral replication factors, such as EBOV L protein and DYNLL1.

To elucidate host factors involved in EBOV replication and transcription, we performed two independent replicates of a genome-wide pooled CRISPR KO screen. For each replicate we transduced the EBOV replicon cell line with the Brunello library and used FACS as a measure of minigenome replication and transcription activity. Cells with significantly decreased eGFP signal over the course of the screen (eGFP-low) were sorted and collected and compared to unsorted cells frozen after library transduction and completion of puromycin selection. The same sequencing and bioinformatics analysis workflow was applied to determine the genes enriched in the eGFP-low fraction over the course of the screen.

Several classes of novel host genes potentially involved in EBOV replication were enriched in the screen (Figure 4C). The top three hits in our replicon screen belong to the protein family of histone modifiers. EHMT1 and EHMT2 are histone lysine methyltransferases ^43^, and EP300 encodes the adenovirus E1A-associated cellular p300 transcriptional co-activator protein that functions as a histone acetyltransferase ^44,45^. We also identified KOs of ubiquitin family proteins (FBXO28, USP7, and USP12) among the enriched genes from the screen. Other host factors that were found to be enriched in the screen included several protein phosphatases/kinases (DUSP6, DUSP18, PRKCI, PPP1R12B, and PPP3CB), immunoglobulin superfamily genes (CD19 and ICAM4), and transcriptional modulators (NRIP1, PSIP1, ZNF865, ZNF182).

### Distinct verified sets of host factor dependencies for replication of diverse viruses identified in genome-wide CRISPR replicon screens

To narrow down hits for follow-up confirmation studies, we identified seven genes recovered independently in the set of genes ranking in the top 1% for each screen replicate (Supplemental Figure 5A and 5B, blue text). Beyond these, we included an additional nine genes ranking in the top 1% in either screen that (a) belonged to similar protein functional groups or (b) corresponded to proteins implicated in EBOV replication in the literature (Supplemental Figure 5B, black text). The selected genes were individually knocked out in the EBOV replicon cell line using the top-performing gRNAs from the Brunello library and then monitored for eGFP expression. Controls included KOs targeting the EBOV L protein, DYNLL1, NPC1, and a cell line harboring a non-targeting gRNA (NT). Independent validation of the decrease in EBOV minigenome eGFP activity was observed for the EHMT1, EHMT2, EP300, and USP7 KO cell lines (Figure 4D).

**Figure 5.**
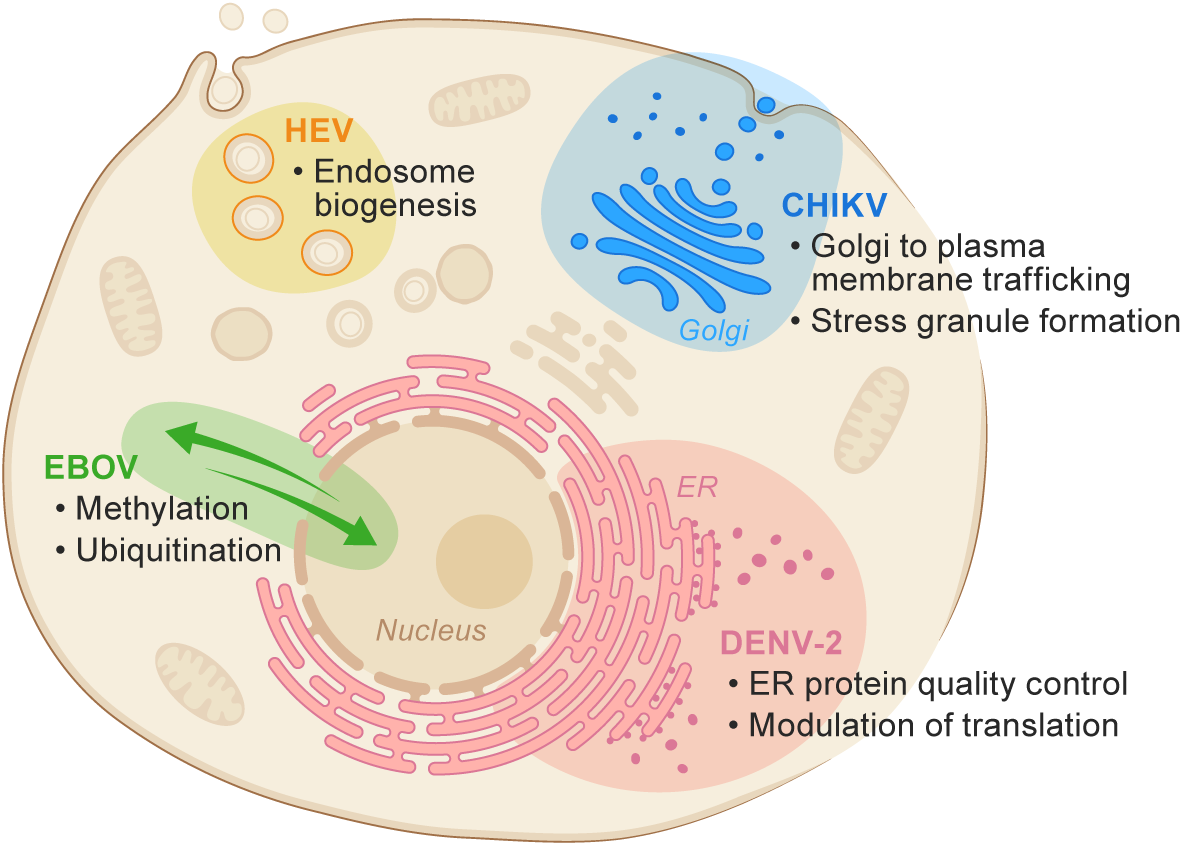
Distinct verified sets of host factor dependencies for replication of diverse viruses identified in genome-wide CRISPR replicon screens. Graphical summary of the host factor dependencies within the cell recovered for DENV-2, CHIKV, and EBOV (this study) as well as hepatitis E virus (HEV) ^56^. Created in https://BioRender.com.

To directly assess the effect on viral replication and transcription among the verified hits, we quantified the EBOV minigenome expression in the EBOV replicon cell lines with the verified KOs ^40^. Depletion of EHMT1, EHMT2, and USP7 resulted in an overall decrease in abundance of all three of the EBOV minigenome RNA products relative to the levels observed in cells transduced with a non-targeting gRNA (Figure 4E; Supplemental Figure 5D).

Because EHMT1 and EHMT2 are known chromatin modifiers, we investigated whether the reduced replicon activity in the EHMT1 and EHMT2 KO cells was due to an indirect effect of suppressing the constitutive expression of the EBOV 4cis integrated in the genome of the EBOV replicon cell line. To test this, we examined the impact of EHMT1 and EHMT2 on transiently transfected EBOV 4cis and minigenome plasmids. Independent EHMT1, EHMT2, and USP7 KOs were generated in the Huh7.5.1–Cas9 cell line (Supplemental Figure 5C). These three KO cell lines plus an Huh7.5.1–Cas9 control cell line harboring a non-targeting gRNA (NT) were co-transfected with an EBOV 4cis plasmid and an EBOV secreted nano-luciferase (SecNLuc) minigenome construct, and SecNLuc activity was assayed 48 h post-transfection. All three KO cell lines had reduced SecNLuc minigenome reporter activity compared to NT control cells (Figure 4F). These data demonstrate that the EBOV replication phenotype detected in EHMT1 and EHMT2 KO cells is not dependent upon EBOV 4cis integration in the genome.

## Discussion

Here we report the development of a genome-wide CRISPR screening approach using viral replicons as the unit of analysis to circumvent limitations of live virus screens and specifically enrich for host factors and pathways involved in the formation and function of viral replication and transcription complexes. A key challenge in developing this approach was the generation of stable viral replicon cell lines. Although viral replicons have been a long-established tool for basic molecular virological analyses and ultra-high-throughput screening campaigns for antiviral compounds, these applications typically are performed in the context of transient transfections and assayed in an arrayed format. In contrast, the pooled format applied for genome-wide CRISPR screening required the development of robust, stable replicon cell lines with >80% fluorescent-positive cells that are responsive to perturbations of viral replication.

In our hands, leveraging an ORF that confers drug resistance via stoichiometric binding to the antibiotic Zeocin was critical for the establishment of the stable DENV-2, CHIKV, and EBOV replicon cell lines. Independent precedence exists for applying the Zeocin gene to dial the dosage of transgene expression in gene delivery vectors ^19^. We speculate that this dose-dependent drug resistance mechanism applies selective pressure to maintain higher replicon RNA copies per cell than drug resistance conferred by an enzyme-mediated drug degradation. The utility of including a PEST degradation tag in the replicon reporter-selection cassette was mixed and had variable effects on reporter stability depending on the specific viral replicon system. Therefore, we found it useful to test a series of eGFP-Zeocin variants in parallel for each new replicon cell line to optimize reporter stability and responsiveness to perturbation before proceeding with the replicon screen.

To test the feasibility of this more targeted CRISPR screening approach, we first focused on developing and testing a DENV-2 replicon cell line. DENV-2 served as an excellent virus to benchmark this approach, because DENV-2 replicon systems are well established, and much is already known about the host factors involved in DENV-2 replication. We determined that a DENV-2 replicon with a Zeocin-eGFP-PEST fusion protein yielded a stable replicon cell line suitable for a pooled FACS-based genome-wide CRISPR screen. The replicon screen enriched all the expected host factors that had been previously identified to be involved in DENV-2 replication via genome-wide screens with live DENV-2. These results establish the feasibility of genome-wide CRISPR KO screens with stable viral replicon cell lines and point to a potentially powerful strategy to functionally screen for host factors and pathways that are specifically required for viral replication.

Beyond DENV-2, generation of stable CHIKV and EBOV replicon cell lines suitable for pooled genome-wide CRISPR screening was feasible, despite their distinct genome organizations and replicon systems. This provided a unique opportunity to begin to investigate the spectrum of host factors and pathways within the cell that are functionally required for viral replication for diverse viral agents. Looking solely at the verified hits for each, distinct patterns of host factor requirements emerged that mirror the compartments each virus uses during infection.

In the follow-up analysis of new hits from the DENV-2 replicon screen, we independently verified new hits via live virus assays. Recovery of deoxyhypusine hydroxylase (DOHH) highlights that modulation of translation initiation factor activity ^46^ may be an additional pathway that specifically influences DENV-2 replication. Indeed, orthogonal studies of DENV-2 translation have implicated a role for this pathway during DENV-2 infection ^47^. Likewise, recovery and validation of zinc finger protein 36, C3H1 type-like 2 (ZF36L2), a factor implicated in attenuating protein synthesis via binding and destabilizing AU-rich RNAs in the cytoplasm ^48^, indicates that pathways influencing RNA stability might be yet another cellular pathway that DENV-2 requires during infection to modulate translation. Taken together, these results corroborate prior data pointing to translation modulation and polyprotein processing in the ER as major cellular pathways required for DENV-2 replication (Figure 5).

The CHIKV and EBOV replicon screens each revealed further distinct sets of host factor dependencies for replication. For CHIKV, recovery of CSDE1, G3BP1, and G3BP2 corroborated and expanded prior targeted studies that have implicated an important role for stress granule components in CHIKV replication ^31,32,35,36^. Separate recovery and validation of GOLGA7 points to a role for Golgi–plasma membrane trafficking and/or the palmitoylation pathways in CHIKV replication complex formation or function. In all, these findings point to CHIKV host factor dependencies spanning the Golgi apparatus, cytoplasm, and plasma membrane (Figure 5).

The verified hits recovered in the EBOV replicon screen (eukaryotic histone methyltransferases 1 and 2 [EHMT1, EHT2, respectively], and ubiquitin specific peptidase 7 [USP7]) are distinct from those recovered in the DENV-2 and CHIKV replicon screens, as well as the hits enriched in prior CRISPR screens with live EBOV ^42,49,50^. However, these genes were intriguing as they have links to prior results from targeted EBOV studies. Recovery of USP7 is consistent with evidence that ubiquitin modification of EBOV proteins NP and VP35 can influence viral replication and transcription ^51,52^. The recovery of EHMT1 and EHMT2 was somewhat paradoxical given that these factors are generally thought to be localized in the nucleus, where they act on histones ^43^, whereas EBOV replication occurs in the cytoplasm. However, both EHMT1 and EHMT2 are also involved in methylation of non-histone proteins ^53^. Moreover, two recent studies have demonstrated a role for EHMT1 and EHMT2 in viral replication of SARS-CoV-2 ^54^ and in the formation and function of viral inclusion bodies during infection with a separate negative-sense RNA virus, Sendai virus ^55^. Taken together, these data suggest a potential role for host factors related to lysine methylation and ubiquitination modifications in EBOV replication complex formation and function (Figure 5). Further biochemical and live virus infection assays are required to clarify these results.

In all, our data provide evidence for the feasibility of using pooled genome-wide CRISPR screens with viral replicons to specifically identify the host factors required for the formation and function of viral replication complexes of diverse RNA viruses. Although we lack a complete understanding of the similarities and differences between stable viral replicon cell lines and cells acutely infected with viruses, the identification of novel host factors that have an impact on replication and transcription when tested in live virus infections indicates that the stable replicon cell lines are recapitulating major aspects of the replication and transcription phase of the viral life cycle. Moreover, the replicon screens with DENV-2, CHIKV, and EBOV replicons have yielded host factors that reflect their distinct biogenesis, localization, and type of viral replication and transcription complexes. Consistent with our results, hits from a separate CRISPR KO screens with a hepatitis E virus (HEV; positive-sense RNA virus family *Hepeviridae*) “suicide replicon” system suggest that another distinct host pathway (early endosomes and the RAB5A gene) is involved in the formation of the HEV replication complex ^56^ (Figure 5).

Combining additional CRISPR screening modalities (CRISPRi/CRISPRa) that facilitate the analysis of essential gene requirements as well as the potential role of gene activation and dosage will likely further enhance the scope of genes detectable by this approach. Likewise, application of this screening approach across diverse RNA viruses—those closely related and more distantly related, including highly virulent viruses that are not easily studied in the live virus—will provide a unique opportunity for comparative analysis of the landscape of host factors/pathways requirements for this phase of the viral life cycle. Such studies offer an opportunity to further elucidate the basic cell biology of the host cell and highlight potential host pathways or gene targets for novel host-directed medical countermeasures.

## Methods

### Cell culture

Grivet (*Chlorocebus aethiops*) kidney epithelial Vero E6 cells were sourced from the American Type Culture Collection (ATCC; #CRL-1586) by BEI Resources and specially banked by Lonza for the Integrated Research Facility at Fort Detrick (IRF-Frederick). Human hepatocyte-derived carcinoma Huh7.5.1 cells were sourced from Apath/Dr. Charles M. Rice and Scripps Research/Dr. Francis V. Chisari. Parental and derivative KO cell lines that were generated at the Chan Zuckerberg Biohub were provided to the IRF-Frederick and grown in Dulbecco’s Modified Eagle Medium (DMEM; Gibco, 11995-040) supplemented with 10% heat-inactivated fetal bovine serum (FBS; Sigma Aldrich, F4135), penicillin-streptomycin, non-essential amino acids, and L-glutamine. Human embryonic kidney (HEK) epithelial 293FT cells used to generate lentiviruses were grown in DMEM supplemented with 10% FBS, penicillin-streptomycin, and L-glutamine. Huh7.5.1 and HEK293FT cells were incubated at 37°C and 5% carbon dioxide (CO2). For knockout (KO) cell lines, media were additionally supplemented with 4 µg/mL puromycin and 100 µg/mL hygromycin. Antibiotics were removed prior to plating the cells for experiments. One day before experiments, 3 x 10^5^ cells per well were seeded into 6-well plates, and 8,000 cells per well were seeded into 384-well plates.

### Virus isolates, reporter viruses, and replicon constructs

#### Viruses

The infectious dengue virus type 2 (DENV-2) clone 16681 was a gift from Jan Carette (Stanford University). This infectious clone was adapted to human chronic myeloid leukemia-derived near haploid (HAP1) cells and contains a mutation leading to a Q399H change in the envelope protein. DENV-2 expressing *Renilla* luciferase (DENV-2-Luc) was also a gift from Jan Carette and expanded on Huh7.5.1 cells. Chikungunya virus (CHIKV) stock IRF0543 (BEI Resources; isolate LR 2006-OPY1) was grown in Vero E6 cells at 37°C and 5% CO2. Cell culture supernatants were collected 3 d post-inoculation. The resulting stocks were aliquoted and stored at -80°C. Viral titers were determined by plaque assay. All work with infectious CHIKV was done in the biosafety level 3 (BSL-3) and biosafety level 4 (BSL-4) containment laboratories at the IRF-Frederick. All the work was approved and done in compliance with the institutional, city, state, and national regulations.

#### DENV-2 replicon constructs

DENV-2 replicon constructs (Supplemental Table 3) were derived from the DENV-2 16681 infectious clone. The region in DENV-2 16681 corresponding to the 5’ UTR to the first *Eco*RI site in the NS1 gene (nucleotides 1–2,876) was replaced with a gene synthesis product (Integrated DNA Technologies) containing *Sac*I–T7–5’UTR–capsid (nucleotides 1–102)–eGFP–Blasticidin–F2A–envelope (last 72 nucleotides)–NS1 (nucleotides 1–455)–*Eco*RI to create the initial eGFP-blasticidin DENV-2 replicon plasmid. All other versions of the DENV-2 replicon with various fluorescent protein-selectable marker cassettes were generated by overlap PCR of the *Sac*I–5’UTR––fluorescent protein–selectable marker cassette–*Bam*HI, then ligated into a *Sac*I/*Bam*HI digested eGFP-blasticidin replicon backbone.

#### CHIKV replicon constructs

CHIKV replicon constructs (Supplemental Table 3) were derived from DNA containing CHIKV^repl^ ^SNAP-nsP3^ ^sg-ZsGreen^, a generous gift from Mark Harris (University of Leeds). The original ZsGreen-puromycin reporter-selection cassette was replaced with a series of eGFP-Zeocin cassettes by digesting the CHIKV^repl^ ^SNAP-nsP3^ ^sg-ZsGreen^ with *Avr*II and *Pme*I and replacing the insert with variants of *Avr*II-eGFP-Zeocin-*Pme*I. SNAP-nsP3 was replaced with wild-type nSP3 by digestion with *Sac*II and *Age*I and replacement with the insert *Sac*II-nsP3-*Age*I. All other eGFP-Zeocin variants were generated in the same way.

#### Ebola virus replicon constructs

The EBOV replicon constructs (Supplemental Table 3) used in this study to generate a stable replicon cell line were derived from gene synthesis of sequences corresponding to isolate Ebola virus/H.sapiens-tc/COD/1976/Yambuku-Mayinga (GenBank accession AF086833.2). Two types of EBOV constructs were developed for generating the stable EBOV minigenome replicon cell line: a PiggyBac donor plasmid for transposon-based integration of the EBOV genes required for replication and transcription into the host genome (pAKFi357), and a separate *in vitro* transcribed RNA encoding the EBOV minigenome negative-sense vRNA (pAKFI368).

### Generation of stable DENV-2 replicon cell line

For *in vitro* generation of DENV-2 replicon RNA, the DENV-2 replicon plasmid was linearized with *Xba*I. First, 1 µg of linearized DENV-2 replicon DNA was transcribed into RNA using a MEGAscript T7 Transcription Kit (Ambion, AM1334) with the addition of 4 mM of m^7^G(5’)ppp(5’)G cap analog (NEB, S1405S) per reaction. The reaction was incubated for 4 h at 33°C and then treated with Turbo DNAse at 37°C for 15 min. Replicon RNA was purified using an RNA Clean and Concentrator-25 kit (Zymo Research, R1017) and stored at -80°C. Huh7.5.1–Cas9 cells were seeded in a 12-well plate (200,000 cells per well) for transfection with purified replicon RNA the following day. Next, 1 µg of purified replicon RNA was combined with an mRNA transfection reagent (Mirus, MIR2225) in accordance with the manufacturer’s protocol. The transfected cells were visualized under a fluorescence EVOS microscope 24 h after transfection for the appearance of eGFP-expressing cells. Cells were placed under antibiotic selection (4 µg/mL blasticidin; 50–200 µg/mL Zeocin) 2 d after RNA transfection. Replicon cells were assayed for eGFP expression by flow cytometry every 2–3 d over the course of 1 mo (Beckman Coulter CytoFLEX) and analyzed using FlowJo software.

### Generation of stable CHIKV replicon cell line

To generate CHIKV replicon RNA, the CHIKV replicon plasmid was linearized with *Not*I. 1 µg of linearized CHIKV replicon DNA was transcribed *in vitro* into RNA using an mMessage mMachine SP6 Kit (AM1340). The reaction was incubated for 4 h at 33°C and then treated with Turbo DNAse at 37°C for 15 min. Replicon RNA was purified using Zymo RNA Clean and Concentrator-25 kit (Zymo Research, R1017) and stored at -80°C. Huh7.5.1.-Cas9 cells were seeded in a 12-well plate (200,000 cells per well) for transfection with purified replicon RNA the following day. 1 µg of purified replicon RNA was combined with Mirus mRNA transfection reagent (Mirus, MIR2225) following the manufacturer’s protocol. The transfected cells were visualized under a fluorescence EVOS microscope 24 h after transfection for the appearance of eGFP-expressing cells. Cells were placed under antibiotic selection (50–200 µg/mL Zeocin) 2 d after RNA transfection. Replicon cells were assayed for eGFP expression by flow cytometry every 2–3 d over the course of a month (Beckman Coulter CytoFLEX) and analyzed using FlowJo software.

### Generation of a stable EBOV replicon cell line

Huh7.5.1–Cas9-P2A-blasticidin cells were plated at a density of 150,000 cells per well and co-transfected with TransIT-X2 reagent (Mirus, MIR6000) + 0.5 μg of a PiggyBac transposase expression plasmid (gift from Hana El Samad, UCSF [current address Altos Labs, Redwood City CA]) and 0.5 μg of a PiggyBac donor plasmid with a hygromycin resistance gene (Addgene, 123943) harboring a multicistronic insert encoding the chicken actin gene (CAG) promoter driving a single ORF consisting of the four EBOV ORFs encoding NP, VP35, VP30, and L protein, separated by P2A ribosome skipping sites, as previously described ^39^ (“EBOV 4cis”). At 24 h post-transfection, cells were transferred to media containing 200 μg/mL hygromycin for a further 10–12 d, with media fresh changes every 2–3 d, to select for a population of cells in which the EBOV 4cis donor plasmid had stably integrated into the host cell genome. After expansion of EBOV 4cis cells surviving selection, aliquots of cells were harvested, lysed with radioimmunoprecipitation assay (RIPA) buffer and tested for the expression of EBOV NP, VP35, and VP30 proteins via western blot analysis with anti-P2A antibodies, and available antibodies raised against EBOV NP, VP35, and VP30 proteins (see western blot methods for details). In parallel, 1 μg of *in vitro* transcribed EBOV minigenome vRNA harboring a Zeocin-eGFP gene fusion was transfected into aliquots of EBOV 4cis cells via chemical transfection (Mirus TransIT mRNA transfection reagent, MIR2256). eGFP signal derived from transcription and replication of the minigenome vRNA in these cells was assessed manually via fluorescence microscopy (Invitrogen EVOS FL imaging system), and the fraction of GFP^+^ cells and signal intensity of eGFP was quantified via flow cytometry (CytoFlex). Within 24–48 h of showing eGFP signal, the EBOV 4cis cells transfected with the minigenome vRNA were transferred to media containing 50 μg/mL Zeocin to select for a stable EBOV replicon cell line that maintained replication and transcription of the minigenome vRNA. A stable population with >90% GFP^+^ emerged after approximately 12 d of growth in media containing 50 μg/mL Zeocin. The stable EBOV replicon cell line was expanded and subjected to increasing concentrations of Zeocin to test growth, GFP signal intensity, and percent GFP^+^ cells. Ultimately, 200–300 μg/mL Zeocin served as the optimal media conditions to maintain the EBOV replicon cells at >95% GFP^+^ cells without negative effects on cell viability.

### Genome-wide CRISPR KO screens

The Human Brunello CRISPR knockout pooled sgRNA library ^23^ was a gift from David Root and John Doench (Addgene, 73178). A lentivirus stock of the library was titered by serial dilution from 1:10–1:1,000 and selected with 4 µg/mL puromycin. A live cell count of each dilution relative to non-transduced and unselected control cells was performed by flow cytometry (Beckman Coulter CytoFLEX) to determine the appropriate dilution for 50% transduction efficiency. One day prior to library transduction, 160 million Huh7.5.1–Cas9-P2A-hygromycin cells (for stable DENV replicon cell line screen) or Huh7.5.1–Cas9-P2A-blasticidin cells (for stable EBOV replicon cell line screen) were split into duplicate sets of T175 flasks and processed independently as replicates for the screen. Replicon cells were transduced with Brunello library lentivirus at 50% transduction efficiency and 500X coverage, grown in 4 µg/mL puromycin for 5–7 d to select for cells harboring integrated gRNA cassettes, and passaged for an additional two weeks. A total of 40 M unselected (“unsorted”) control replicon library cells were collected 6 d after library transduction. On Day 20, after library transduction, each screen replicate was separately pooled and sorted based on eGFP fluorescence using a SONY SH800 FACS. The bottom 20% (“eGFP-low”) and top 10% (“eGFP-high”) of eGFP-expressing cells were separately sorted into collection tubes, washed in phosphate buffered saline (PBS), pelleted by centrifugation, and stored at -20°C. A total input of 40–60 M replicon library cells were FACS-sorted per replicate to obtain >500X library coverage.

Genomic DNA was extracted from the cell pellets of the sorted cells using the Quick-DNA Miniprep Plus kit (Zymo Research, D4069) and from cell pellets of the unsorted cells using the Qiagen Blood Maxi Kit (Qiagen, 51194). The genomic DNA preps were further purified from contaminants (Zymo Research OneStep PCR Inhibitor Removal Kit, D6030). Sequencing libraries were prepared from extracted genomic DNA by PCR amplification using barcoded Illumina P7 and P5 primers of the gRNA cassettes. The entire prep of extracted gDNA was barcoded and amplified (1 µg gDNA/ 50 µl PCR reaction) for each screen condition. Parallel amplified libraries were pooled and purified by SPRIselect beads (Beckman Coulter, B23318) and library size, and concentration were quantified via the Agilent TapeStation. Barcoded libraries were pooled and sequenced for 20 cycles on an Illumina NextSeq 550 High Output run, with a custom sequencing primer that allows for sequencing to begin at the first base of the gRNA cassette.

### Gene enrichment analysis and hit calling

Demultiplexed FASTQ files were analyzed using the ’count’ subcommand of MAGeCK software (v0.5.9.4) ^24^ to quantify gRNA abundance by matching reads to the gRNA library sequences, with the normalization method set to ’total’. The gRNA count tables were subsequently analyzed using the ‘test’ subcommand (RRA) of MAGeCK software v0.5.9.4 to calculate positive enrichment scores for each gene. The two-step process was automated using Nextflow v21.10.6. The analysis workflow standardizes metadata conventions. It is available at https://github.com/czbiohub-sf/CRISPRflow/blob/main/modules.nf.

The gene enrichment score is calculated by taking the negative base-10 logarithm of the calculated positive robust rank aggregation (RRA) score. For hit calling, genes that ranked in the top 1% with a log fold change (LFC) >0 from MAGeCK RRA analysis (Supplemental Table 1) were broadly considered for follow-up validation studies. For the DENV-2 replicon screen, we compared the top 1% gene candidate list to gene lists from previous genome-wide DENV-2 CRISPR screens ^7,8,10^. We selected 15 genes for follow-up validation that, to our knowledge, had not been identified in prior genome-wide CRISPR screens. For the CHIKV replicon screen, we verified the top 20 genes from the top 1% gene candidate list. These included 18 genes that, to our knowledge, had not been identified in prior genome-wide CRISPR screens and 2 genes (positive controls) that are known CHIKV host factors: G3BP1 and G3BP2. For the EBOV replicon screen, we focused on the top 1% of hits from two independent biological replicates of EBOV replicon screen. From these, we selected seven hits that consistently showed up in the two independent replicates of EBOV screens. Moreover, based on literature, we selected nine more genes that showed up in the top 1% of any one of our screen replicates that could be involved in EBOV replication. DYNLL1 was included as a positive control since it has been previously implicated in EBOV replication ^41^.

### Generation of KO cell pools

A set of gRNAs targeting each candidate gene from the replicon CRISPR screens were selected from the Brunello library (Supplemental Table 2) and cloned into pLentiGuide-puro plasmid and packaged into lentivirus particles using HEK293FT cells. Huh7.5.1–Cas9-P2A-hygromycin cells were seeded at 80,000 cells per well in 24-well dishes. The following day, each well was transduced with lentiviruses expressing the gRNA cassette for gene KO. Cells were selected for gRNA cassette integration with 4 µg/mL puromycin 2 d post-transduction, passaged for a minimum of 7 d, and either harvested for genotyping or downstream cell-based assays.

### Genotyping CRISPR editing of KO cell pools

CRISPR editing in the KO cell pools was performed for the gene KOs that influenced viral replication. First, genotyping primers targeting a 300-bp region around the predicted genomic cut site were designed for each gRNA using the open-source software program protospaceJAM (https://protospacejam.czbiohub.org/). gDNA was extracted from the KO cell pools and unedited Huh7.5.1 cells using the Zymo Quick-DNA Plus Miniprep kit (Zymo Research, D4069) and then used as the template for genomic PCRs. Purified PCR products were submitted for Sanger sequencing and analyzed using Synthego ICE analysis software (https://ice.synthego.com/).

### Arrayed KO for validation of CRISPR screen hits

The CRISPR screen hits were verified by arrayed KO assay in which selected hits from the CRISPR screen were individually knocked out in replicon cells using the top-performing gene-specific gRNAs from the Brunello library. Top-performing gRNAs were cloned in a lentivirus expression construct (pLentiGuide-puro) and lentivirus particles were produced using HEK293FT cells. Briefly, the 10,000 replicon cells were plated in 96-well plates and transduced with lentiviruses expressing the gRNAs targeting specific genes to be knocked out in triplicate. The next day, media were replaced with puromycin selection media (4 μg/mL) to select lentivirus-transduced cells. After all cells in the negative control samples died, the selection media were replaced with regular media and CHIKV minigenome activity was recorded every 3–4 d for the duration of arrayed KO experiment (20–25 d), using CytoFLEX to measure eGFP signal.

### Western blots

Protein expression for viral and host protein expression was analyzed by western blots. Briefly, 1–2 million cells were trypsinized and pelleted. After one PBS wash, the pellets were frozen at - 80°C. At the time of lysing the cells, pellets were thawed on ice and lysed in ice cold RIPA lysis buffer (Thermo Fisher Scientific, 89901) containing 1X phosphatase/ protease inhibitor cocktail (Sigma Aldrich, 11836153001) by vortexing every 5 min for 30 min. Lysates were then centrifuged at 14.000 x*g* for 10 min. Supernatant was collected in a fresh tube and a 4X Laemmli buffer was then added, followed by boiling the samples at 95°C for 10 min. Proteins were then separated on 4–15% sodium dodecyl sulfate-polyacrylamide gel electrophoresis (SDS-PAGE) tris-glycine buffered gels (Thermo Fisher Scientific, 4–15% Mini-PROTEAN TGX Precast Protein Gels, 4561085) and transferred to nitrocellulose membranes using the Biorad Trans-Blot turbo transfer system (Bio-Rad laboratories) in accordance with the manufacturer’s recommended protocol. Primary antibodies used were as follows - DENV NS2B (Genetex, GTX124246), DENV NS3 (Genetex, GTX629477), DENV NS4B (GeneTex, GTX103349), GFP (GeneTex, GTX113617), vinculin 7F9 (Santa Cruz Biotechnology, sc-73614). CHIKV nsP2 (Genetex, GTX135188), CHIKV nsP3 (Genetex, GTX135189), CHIKV nsP4 (Thermo Fisher Scientific, PA5-117443), beta tubulin (Cell Signaling, 15115) NP (IBT Bioservices, 0301-012), VP35 (Kerafast, Kf Ab02366-1.1), VP30 (GeneTex, GTX134035), 2A (Novus, NBP2-59627),EHMT1 (Abcam, ab241306), EHMT2 (Thermo Fisher Scientific, MA5-14880), USP7 (Thermo Fisher Scientific, PA5-34911), and beta tubulin (Thermo Fisher Scientific, MA5-16308).

For experiments involving live CHIKV infections, cell lysates were prepared using 4X SDS buffer (Thermo Fisher Scientific, 50-198-643), supplemented with a protease inhibitor cocktail (Sigma Aldrich, 11836153001) at a 1X final concentration. To prepare cell lysates, media were removed from the well and 500 µL of lysis buffer per well were added to the cell monolayers. For supernatant, 334 µL of SDS buffer was mixed with 1 mL of supernatant. Subsequently, 3 µL of the lysate were analyzed using the Jess Automated Western Blot System (ProteinSimple), following the manufacturer’s recommended protocol. Presence of E1 protein was detected using an anti CHIKV E1 [HL2069] antibody (GeneTex, GTX637973), used at a dilution of 1:500 followed by an Anti-Rabbit Secondary HRP Antibody (Bio-Techne, 042-206).

### Live virus infection assays

#### DENV-2

Huh7.5.1 cells (and derivative KO cell pools) were plated in triplicate per condition at 8,000 cells/well in 96-well plates and infected with either DENV-2 16681 or DENV-2-Luc reporter virus at MOI = 0.1. For infection with DENV-2 16681 virus, cells were collected for analysis 48 h after exposure to DENV-2 16881 virus and analyzed by RT-qPCR. For infection with DENV-2-Luc reporter virus, cells were collected for luminescence analysis 48 h after exposure.

#### CHIKV

Virus was obtained through BEI Resources, NIAID, NIH, as part of the WRCEVA program: isolate LR 2006-OPY1 (product id, NR-49741). Cells were infected with CHIKV at an MOI = 0.01 or 0.1. Briefly, the virus was diluted to the desired MOI in an inoculation volume of 0.5 mL. This inoculum was then added to the cells and incubated at 37°C. After 1 h, the inoculum was removed, and cells were washed once with PBS (Thermo Fisher Scientific, 10-010-023). The cells were then overlaid with 2 mL of fresh DMEM supplemented with 10% FBS. Cells and supernatants were collected at 7, 11, 18, 24, and 32 h post-infection. Samples were processed as follows: (i) for RNA extraction, cells were treated with TRIzol LS, and (ii) for protein analysis, cells were lysed in 4X SDS buffer for subsequent western blot analysis. RNA was extracted from the TRIzol LS-treated samples, and vRNA levels were quantified using RT-qPCR.

### RT–qPCR quantitative analysis of vRNA

#### DENV-2

Cell lysates harvested from DENV-infected and control samples were analyzed by RT-qPCR using the Cells-to-Ct kit (Thermo Fisher Scientific, A25600) in accordance with manufacturer’s protocol for qPCR readout of vRNA relative to 18S RNA (housekeeping gene). The following qPCR primers were used: universal DENV forward, 5’-GGTTAGAGGAGACCCCTCCC-3’; universal DENV reverse, 5’-GGTCTCCTCTAACCTCTAGTCC-3’; 18S forward, 5’-AGAAACGGCTACCACATCCA-3’; and 18S reverse: 5’-CACCAGACTTGCCCTCCA-3’. Raw Ct values were extracted and normalized to 18S for further analysis.

#### CHIKV

RNA was extracted from cells or culture supernatants treated with TRIzol LS (Thermo Fisher Scientific, 10296028) using the Maxwell RSC Viral Total Nucleic Acid Purification Kit (Promega, AS1330) and the automated Maxwell RSC extractor (Promega, AS4500). Briefly, 200 µL of TRIzol LS-treated sample were added to 200 µL of the kit’s lysis buffer and vortexed for 10 s. The entire mixture was then transferred to the extraction kit cartridge and processed on the extractor using the proprietary method specified for the extraction kit. The sample was eluted in 50 µL of nuclease-free water, and RNA was quantified using a Nanodrop 8000 spectrophotometer (Thermo Fisher Scientific). All RNA samples were stored at -80°C until use. vRNA was quantified using RT-qPCR targeting the GP2 gene of CHIKV, according to a previously described protocol ^57^. Briefly, the RT-qPCR reactions were setup using the TaqPath 1-Step RT-qPCR Master Mix, CG, (Thermo Fisher Scientific, A15299) on a QuantStudio 7 Pro system (Applied Biosystems, A43183). The cycling conditions, which included reverse transcription, were as follows: 25°C for 2 min, 50°C for 15 min, and 95°C for 2 min, followed by 40 cycles of denaturation at 95°C for 3 s and amplification at 60°C for 30 s. The primers and synthetic sequence copy standard (gBlock) were designed in house and synthesized by Integrated DNA Technologies, Coralville, IA, USA, with the following sequences: forward primer, 5’-AAGCTCCGCGTCCTTTACCAAG-3’; reverse primer, 5’-CCAAATTGTCCTGGTCTTCCT-3’; probe, FAM-CCAATGTCTTCAGCCTGGACACCTTT-BHQ1. Viral cDNA copies were quantified using a synthetic gBlock that had been serially diluted tenfold from 10 million copies to one copy to create a standard curve. The sequence of this gBlock was 5’-ctcataccgcatccgcatcagctaagctccgcgtcctttaccaaggaaataacatcactgtaactgcctatgcaaacggcgaccatgccgtcacagttaaggacgccaaattcattgtggggccaatgtcttcagcctggacaccttttgacaacaaaatcgtggtgtacaaaggtgacgtttacaacatggactacccgccctttggcgcaggaagaccaggacaatttggcgata-3’.

#### EBOV

A strand-specific RT-qPCR assay was used to identify and measure levels of 3 RNA species in replicon cells: vRNA, cRNA, and mRNA. Cells were collected at the end of the arrayed KO experiment by trypsinizing and pelleting them. Cell pellets were frozen at -80°C until ready for RNA extraction. RNA was extracted using the Zymo Quick-RNA Miniprep Plus Kit (Zymo Research, R1058), quantified using nanodrop and aliquoted to freeze at -20°C. Quality of extracted RNA was determined using the Agilent Tapestation. Synthesis of cDNA was performed using LunaScript RT supermix kits. The LunaScript RT supermix (primer-free) kit (NEB E3025S) was used to synthesize cDNA for vRNA, cRNA, and mRNA; the LunaScript RT supermix kit (NEB E3010S) was used to synthesize cDNA for the glyceraldehyde-3-phosphate dehydrogenase (GAPDH) mRNA control. RT reaction mixtures were prepared using 1 μg RNA/reaction following the manufacturer’s instructions. Each reaction was performed in triplicate. Strand-specific primers used for RT are as follows: mRNA (5’-CCAGATCGTTCGAGTCGTtttttttttttttttttttvn-3’), cRNA (5’-GCTAGCTTCAGCTAGGCATCacacaaagattaaggctatcaccg-3’), vRNA (5’-GGCCGTCATGGTGGCGAATagctttaacgaaaggtctgggctc-3’), GAPDH (random hexamers +dt primers in Lunascript RT Supermix kit). These primers specifically target the RNA species of interest and incorporate a specific barcode sequence (in capital letters) which is then used for amplification during the next qPCR step. For each qPCR, 1 μL of cDNA was used as input for qPCR reactions using Luna Universal qPCR Master Mix (NEB M3003S) following the manufacturer’s instructions. Primers used for qPCR: vRNA (forward, 5’-GGCCGTCATGGTGGCGAAT-3’; reverse, 5’-ACACAAAGATTAAGGCTATCACCG-3’), mRNA (forward, 5’-CTACCTGAGCACCCAGTCCG-3’; reverse, 5’-CCAGATCGTTCGAGTCGT-3’), cRNA (forward, 5’-AGCTTTAACGAAAGGTCTGGGCTC-3’; reverse, 5’-GCTAGCTTCAGCTAGGCATC-3’), and GAPDH (forward, 5’-AAGGGCCCTGACAACTCTTTT-3’; reverse, 5’-CTGGTGGTCCAGGGGTCTTA-3’). Raw Ct values were extracted and normalized to GAPDH for further analysis.

### Luciferase assays

Cells infected with DENV-2-Luc were first stained with Hoechst for 5 min in a 96-well glass-bottom plate as a proxy for cell number (used for normalization). Next, 405 nm fluorescence per well was measured on a Spectramax i3x plate reader (Molecular Devices). Cells were washed three times with PBS and lysed. Then *Renilla* luciferase assay reagent was added to each well of the Promega *Renilla* luciferase assay reagent (E2810) in accordance with the manufacturer’s protocol. Luciferase signal was measured using luminescence readout per well on the Spectramax i3x plate reader. Luminescence measurements were normalized to Hoechst fluorescence for each well.

For EBOV replicon experiments, a luciferase assay was performed using Promega Nano-Glo Luciferase Assay kit (Promega, N1120) following the manufacturer’s instructions. Briefly, 48 h post-transfection using Mirus TransIT-LT1 DNA transfection kit (Mirus, MIR2300), plates were removed from the incubator and culture media were mixed by gently shaking the plates before taking aliquots for measuring secreted nano-luciferase (SecNLuc) activity. The culture media and Nano-Glo reagent were mixed well in a 1:1 ratio. After 3–5 min, luciferase signal was measured using an Envision plate reader (Perkin Elmer). Cell counts were performed by flow cytometry (Beckman Coulter, CytoFLEX) to normalize luciferase activity.

## Data accessibility

Data are deposited in the NCBI Gene Expression Omnibus (GEO) repository and short read archive (SRA). They can be accessed via GEO study id GSE284379 and SRA project id PRJNA1198907.

## Supplemental Files

**SupplementalFile1.pdf**, Supplemental Figures.

**Supplemental Table 1_MAGeCK-RRA.xlsx**, Gene enrichment summary statistics output from MAGeCK analysis run for each screen.

**Supplemental Table 2_gRNA sequence.xlsx**, gRNA target sequences utilized in this study for specific KO cell line generation.

**Supplemental Table 3_PlasmidConstructs.xlsx**, list of plasmid constructs used in this study.

## Acknowledgements

We thank the CZB-SF Genomics platform lead Norma Neff and team members Angela Detweiler, Honey Mekonen, Sheryl Paul, and Amanda Seng for consultation and technical support for library preparation and sequencing throughout this study; and Sandy Schmid (CZB-SF) for feedback over the course of these experiments and critical reading of the manuscript. We also thank Anya Crane and Jiro Wada (Integrated Research Facility at Fort Detrick, Division of Clinical Research, National Institute of Allergy and Infectious Diseases, National Institutes of Health, Frederick, MD, USA) for critically editing the manuscript and figure preparation, respectively, and the Cell Culture staff of the Integrated Research Facility at Fort Detrick for providing cells.

The CRISPR screens and follow-up validation analyses described in this study were funded by the Chan Zuckerberg Initiative. The live chikungunya virus infection experiments were performed at the Integrated Research Facility at Fort Detrick, Division of Clinical Research, National Institute of Allergy and Infectious Diseases, National Institutes of Health, Fort Detrick, Frederick, MD, and were supported in part through Laulima Government Solutions, LLC, prime contract with the U.S. National Institute of Allergy and Infectious Diseases (NIAID) under Contract No. HHSN272201800013C (S.L., A.M.). J.H.K. performed this work as an employee of Tunnell Government Services (TGS), a subcontractor of Laulima Government Solutions, LLC, under Contract No. HHSN272201800013C.

The views and conclusions contained in this document are those of the authors and should not be interpreted as necessarily representing the official policies, either expressed or implied, of the U.S. Department of Health and Human Services, the U.S. Department of Defense, and U.S.

Department of the Army, or of the institutions and companies affiliated with the authors, nor does mention of trade names, commercial products, or organizations imply endorsement by the U.S. Government.

