## Supplemental File 1 for "Genome-wide CRISPR knockout screening with viral replicons for identification of host factors involved in viral replication"

### Supplemental File 1. Cheng, K et al., Genome-wide CRISPR knockout screening with viral replicons for identification of host factors involved in viral replication

#### Supplemental Figures

Supplemental Figure 1. Generation of a stable and responsive dengue virus type 2 replicon reporter cell line suitable for genome-wide CRISPR knockout screening: Replicon construct design and expression of viral proteins.

A, Schematic diagram of the dengue virus type 2 (DENV-2) 16681 genome. B, Schematic diagram of the DENV-2 16681 infectious clone. Structural genes were replaced with a fluorescent protein-selectable marker reporter cassette. Reporter cassettes: eGFP-blasticidin, eGFP-Zeocin, Zeocin-eGFP-PEST. C, Western blot confirmation of expression of DENV-2 proteins NS2B, NS3, NS4B, and eGFP reporter in the Huh7.5.1-Cas9 DENV-2 replicon cell line compared to the parental Huh7.5.1–Cas9 cell line. Loading control: vinculin.

Supplemental Figure 2. CRISPR replicon screen identifies known and novel host factors involved in dengue virus type 2 translation and replication: Genetic confirmation of independently generated gene knockouts.

A, Sanger sequencing traces of target regions PCR amplified from genomic DNA extracted from wildtype and knockout (KO) cell line populations.

Supplemental Figure 3. Genome-wide CRISPR knockout screen with a stable chikungunya virus replicon cell line: Chikungunya replicon design overview, summary of screen hits, and genetic validation of knockout cell lines.

A, (CHIKV) genome schematic (top) and diagram of previously established nsP3-SNAP-tagged zsGreen-puromycin CHIKV replicon [31] and the replicon variants generated and tested in this study. B, Western blot confirmation of expression of CHIKV nonstructural proteins nSP2, nSP3, and nSP4 and the eGFP reporter in the Huh7.5.1–Cas9 CHIKV replicon cell line compared to the parental Huh7.5.1–Cas9 cell line. Loading control: tubulin. C, Genetic confirmation of gene targeting in independently generated populations of Huh7.5.1 knockout (KO) cell lines. Sanger sequencing traces of target regions PCR amplified from genomic DNA extracted from wildtype and KO cell line populations.

Supplemental Figure 4. Genome-wide CRISPR knockout screen with an Ebola virus replicon cell line: Minigenome replicon system overview

A, Schematic of Ebola virus (EBOV) genome structure. B, Schematic of EBOV minigenome replicon used in this study. C, Western blot representing expression of 4cis proteins in the replicon cells. The cell lysates for replicon cells and control cells were probed with anti-NP, anti-VP35 and anti-VP30 antibodies. Parental Huh7.5.1-Cas9 (WT) cells were analyzed in parallel; loading control: tubulin. D, Western blot with P2A antibodies to assess expression and processing of the 4cis proteins harboring a P2A tag (NP, VP35, and VP30, and the integrated Cas9 gene) present in the parental Huh7.5.1–Cas9 cell line and the stable Huh7.5.1-Cas9 EBOV replicon cell line; loading control: tubulin. E, Relative expression levels of the three different types of minigenome RNA expected in the replicon cell line. Total cellular RNA was isolated from replicon cells and used as an input for a strand-specific reverse transcription quantitative PCR (RT-qPCR) assay designed to detect the negative-sense minigenome viral RNA (vRNA), the positive-sense RNA transcript complementary to the vRNA that is generated during viral replication (cRNA), and the positive-sense messenger RNA (mRNA) that is transcribed from the minigenome vRNA template. The values plotted here are fold expression changes of vRNA, mRNA and cRNA relative to the housekeeping control gene glyceraldehyde-3-phosphate dehydrogenase (GAPDH).

Supplemental Figure 5. Genome-wide CRISPR knockout screen with an Ebola virus replicon cell line: Selection and validation of hits.

A, Correlation of hits across the two independent biological replicates of the screen. B, List of hits selected for validation using an arrayed knockout (KO) assay. Blue text indicates common genes enriched across the two independent biological replicates. Black text indicates genes among the top 200 enriched hits in either replicate of the screen that, based on literature survey, have been implicated to play a possible role in EBOV replication and transcription. C, Western blot validation of depletion of host factors EHMT1, EHMT2, and USP7 in the KO cell lines. Cell pellets collected from KO cells were lysed and probed with specific antibodies to assess protein levels in parental and KO replicon cell lines, with tubulin serving as a loading control. D, Relative ratios of viral RNA (vRNA), messenger RNA (mRNA), and RNA complementary to vRNA (cRNA) in the different knockout cell lines. Fold expression changes in vRNA, mRNA and cRNA relative to housekeeping control gene glyceraldehyde phosphate dehydrogenase (GAPDH). Values plotted correspond to mean ± SD for three independent technical replicates.

Supplemental Figure 1

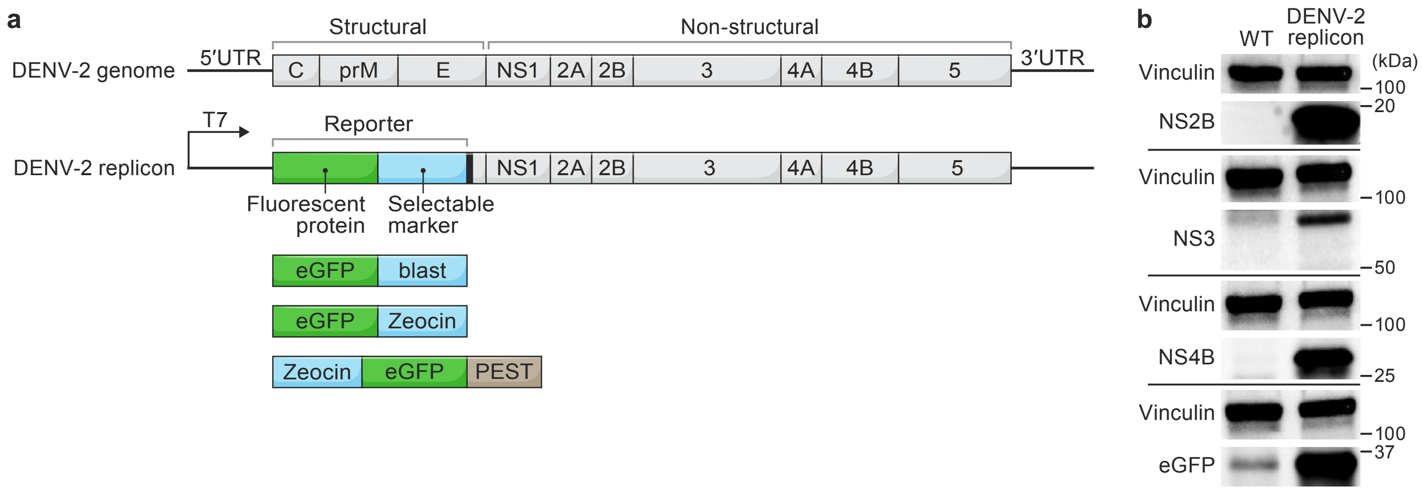

Supplemental Figure 2

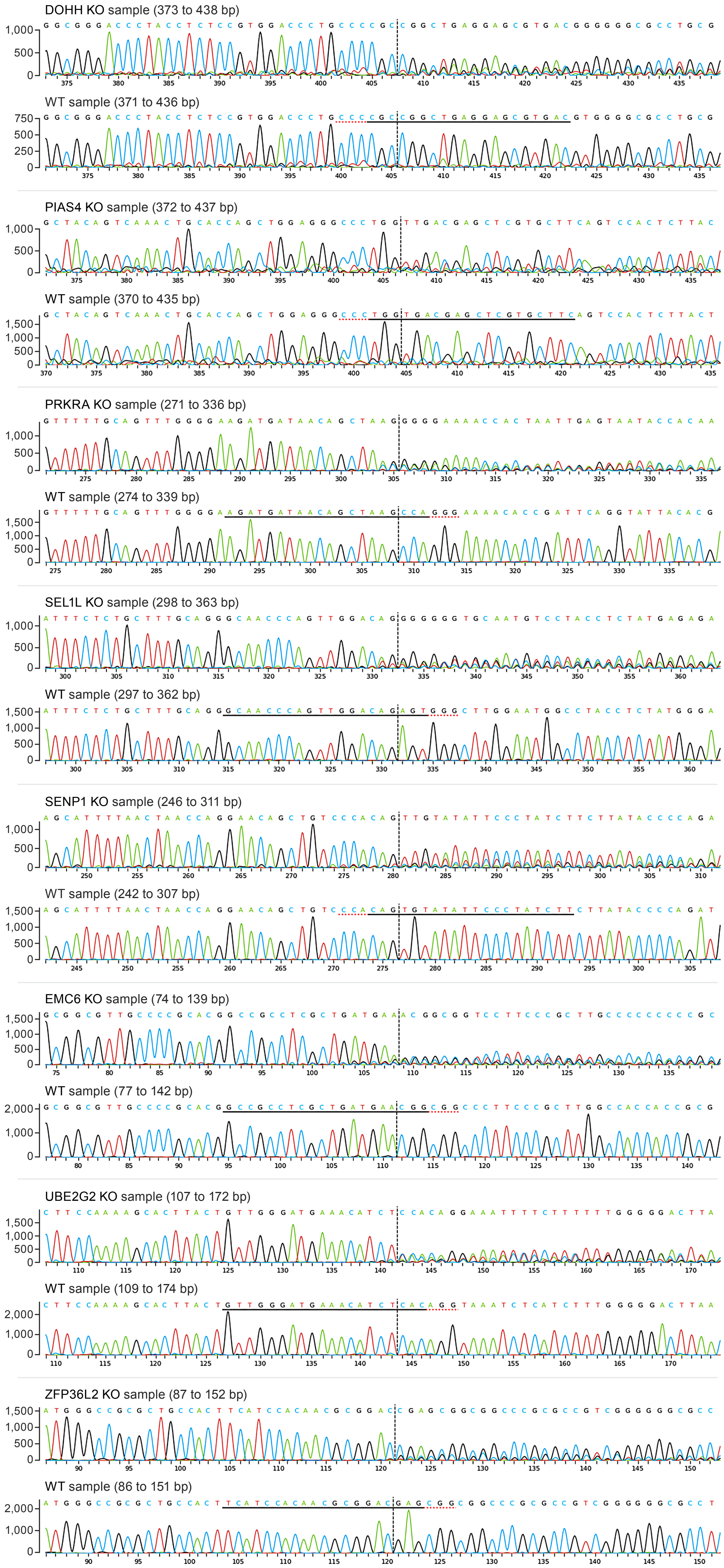

### Supplemental Figure 3

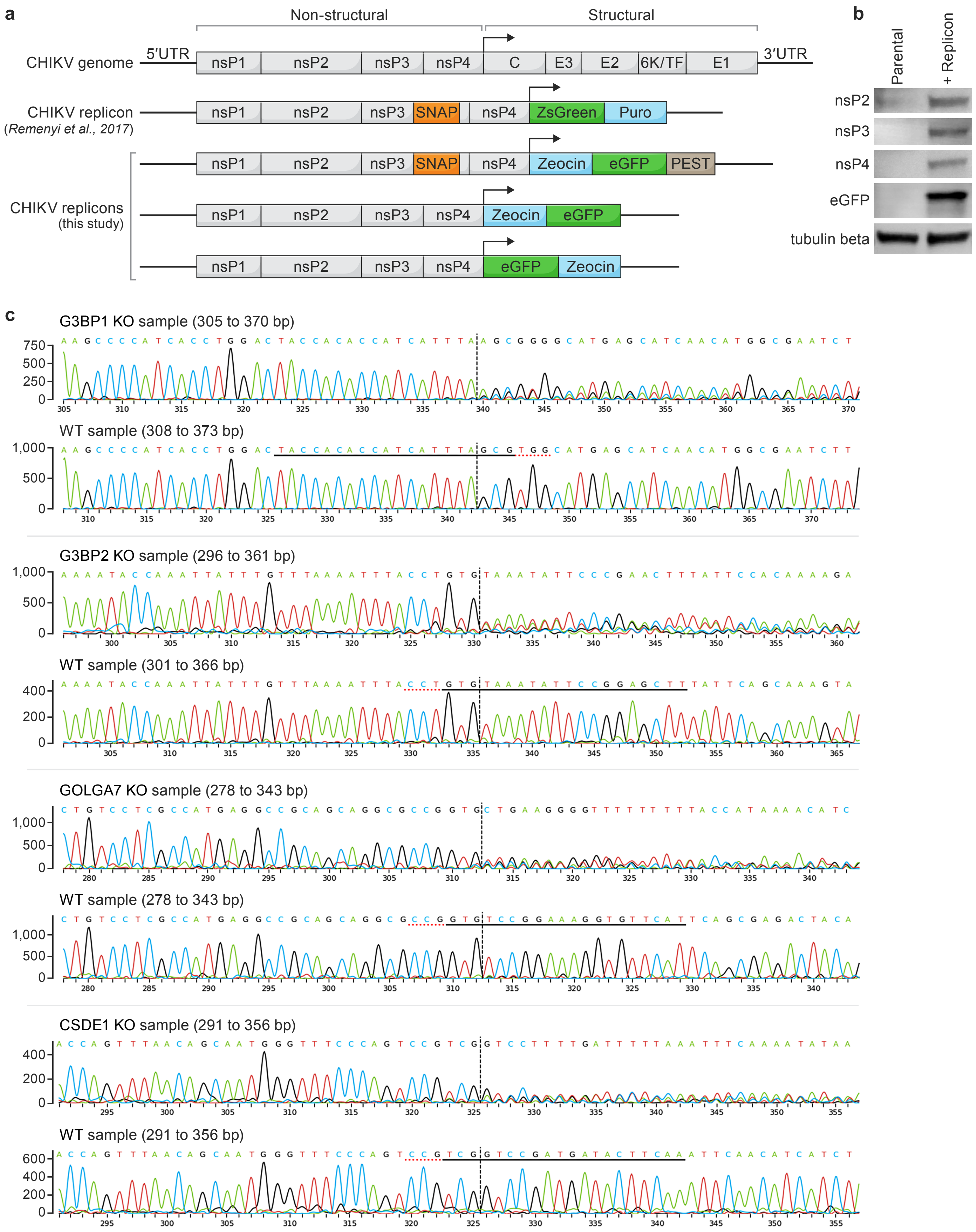

### Supplemental Figure 4

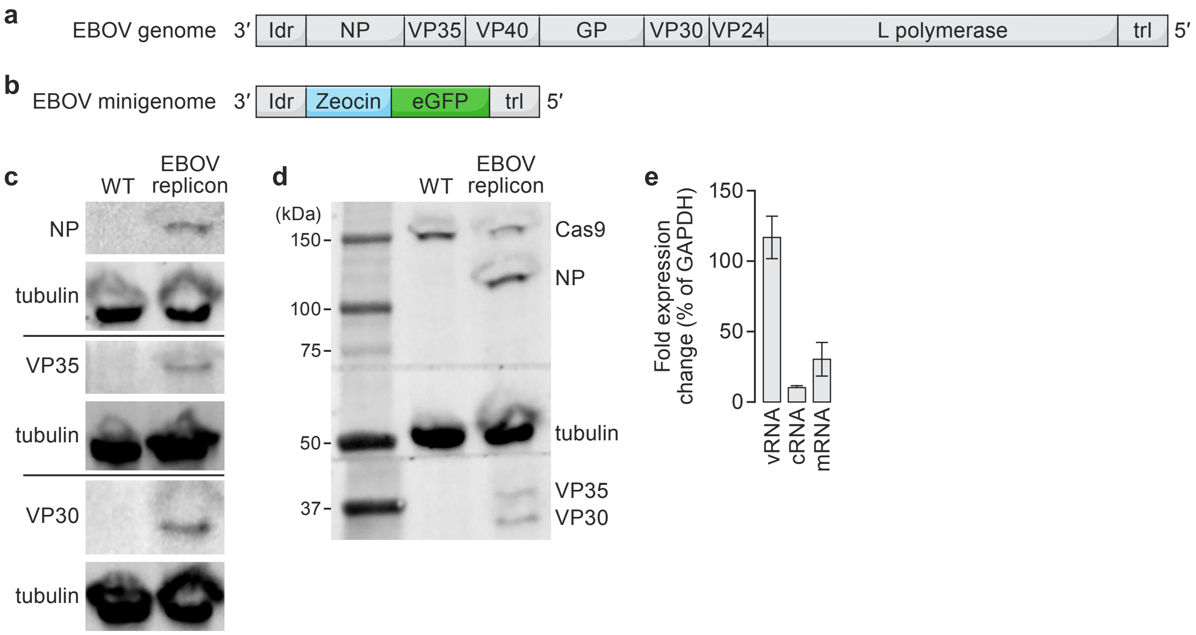

### Supplemental Figure 5

a

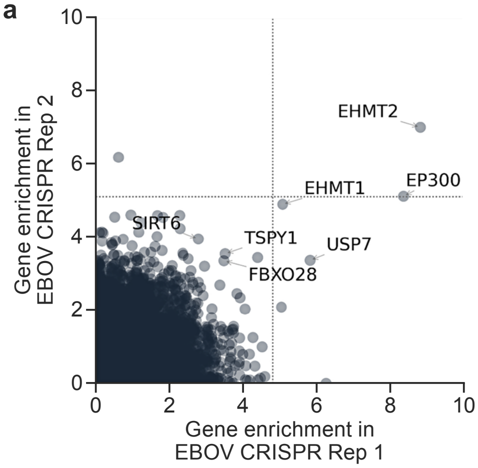


b

Hits selected for validation by arrayed KO assay

| | |
|---|---|
| EHMT1 | GALNT7 |
| EHMT2 | MIR1-1HG |
| EP300 | PPP1CC |
| USP7 | PPP1R12B |
| FBXO28 | PPP3CB |
| SIRT6 | PRKCI |
| TSPY1 | PRKRA |
| DYNLL1 | SUMO3 |
| TP53BP2 | |

c

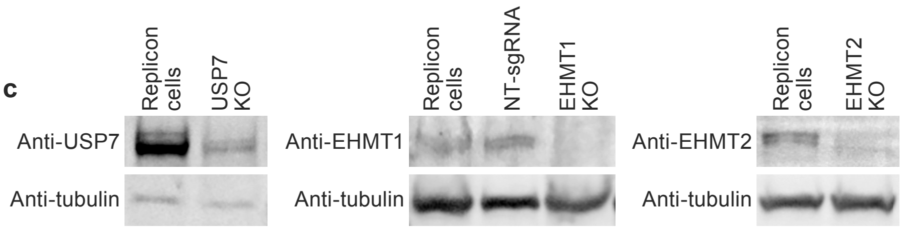


d

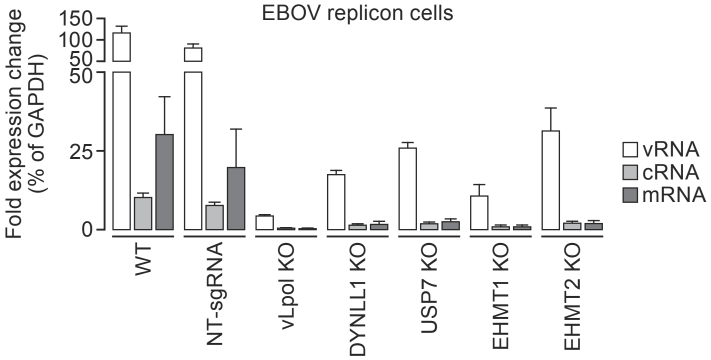
